# Infection of immune competent macrophages expressing functional Slc11a1 alters global gene expression, regulation of metal ions, and infection outcomes

**DOI:** 10.1101/2021.03.18.436026

**Authors:** Lara N. Janiszewski, Michael Minson, Mary A. Allen, Robin D. Dowell, Amy E Palmer

## Abstract

Nutritional immunity involves cellular and physiological responses to invading pathogens, such as limiting iron availability, increasing exposure to bactericidal copper, and manipulating zinc to restrict the growth of pathogens. Manipulation of zinc at the host-pathogen interface depends on both the pathogen’s identity and the nature of the host cell. Here we examine infection of bone marrow-derived macrophages from 129S6/SvEvTac mice by *Salmonella* Typhimurium. Unlike Balb/c and C57BL/6 mice, 129S6/SvEvTac mice possess a functional Slc11a1 (Nramp-1), a phagosomal transporter of divalent cations. We carried out global RNA sequencing upon treatment with live or heat-killed *Salmonella* at 2 Hrs and 18 Hrs post-infection and observed widespread changes in metal transport, metal-dependent, and metal homeostasis genes, suggesting significant remodeling of iron, copper, and zinc availability by host cells. Changes in host cell gene expression suggest infection increases cytosolic zinc while simultaneously limiting zinc within the phagosome. Using a genetically encoded sensor, we demonstrate that cytosolic labile zinc increases 36-fold 12 hrs post-infection. Further, manipulation of zinc in the media alters bacterial clearance and replication, with zinc depletion inhibiting both processes. Comparing our results to published data on infection of C57BL/6 macrophages revealed notable differences in metal regulation and the global immune response, with 129S6 macrophages transitioning from M1 to M2 polarization over the course of infection and showing signs of recovery. Our results reveal that functional Slc11a1 profoundly affects the transcriptional landscape upon infection. Further, our results indicate that manipulation of zinc at the host-pathogen interface is more nuanced than that of iron or copper. 129S6 macrophage leverage intricate means of manipulating zinc availability and distribution to limit the pathogen’s access to zinc while simultaneously ensuring sufficient zinc to support the immune response.

**Author summary:** Metal ions play an important role in influencing how immune cells such as macrophages respond to infection by pathogens. Because metal ions are both essential to survival, as well toxic when present is excessive amounts, the host and the pathogen have evolved diverse strategies to regulate metal acquisition and availability. Here, we show that the metal transporter slc11a1 plays a critical role in defining the host response to Salmonella infection. Infection causes widespread changes in expression of metal regulatory genes to limit the pathogen’s access to iron, increase its exposure to copper, and remodel zinc to ensure increased zinc in the cytosol and limited zinc for the pathogen. Macrophages expressing functional slc11a1 have a different profile of metal regulation and vastly different outcomes compared to immune compromised macrophage, demonstrating significantly different nutritional immune responses in immune competent versus immune compromised macrophages.

## Introduction

Pathogenic organisms must acquire essential micronutrients such as iron, zinc, and manganese to sustain growth and maintain pathogenicity. To protect itself, a host organism in turn attempts to sequester these nutrients, as unfettered access can allow for the exponential growth of an invasive microbe population(1). Metal ion sequestration occurs on a systemic level, evidenced by the sharp decrease of free metal ions in serum upon infection(2). Additionally, phagocytic immune cells manipulate the intracellular localization of metal ions to limit the viability of engulfed pathogens, but this process is more complex than unilateral sequestration. Within macrophages, iron (Fe) ions are bound by cytosolic storage proteins and transported out of phagosomal compartments(3), while these same compartments are flooded with bactericidal copper (Cu) ions(4, 5). The role of zinc (Zn) in nutritional immunity in macrophages is more nuanced than that of Fe or Cu, as it can be used either to poison or starve a pathogen(1). Some microbes have developed strategies to counter these tactics, including siderophores to scavenge Fe ions, oxidases to detoxify Cu, exporters to expunge excess metals, and high-affinity importers to overcome metal limitation(1,2,6). There is a growing body of work suggesting that both the pathogen and the host have mechanisms to regulate Zn accessibility(1, 7), however the overall picture of Zn regulation during infection is not well understood.

Zn is an essential micronutrient required for growth, proliferation, and several fundamental biological processes. In mammals, Zn is a requisite cofactor either structurally or catalytically for approximately 10% of the proteome(8), including many transcription factors and enzymes. As with other metal ions, excess Zn can be toxic, and cells tightly regulate its availability. Mammalian cells contain hundreds of micromolar total Zn but most of this is bound to proteins, enzymes and other ligands such that labile (or exchangeable) Zn, as measured by fluorescent sensors, is typically in the hundreds of picomolar range in the cytosol(9–11). There are fewer tools for, and hence less consensus on, the concentration of labile Zn in organelles. However, multiple different fluorescent sensor platforms have suggested that labile Zn distribution in organelles is heterogeneous(11–15). Zn is regulated by membrane-specific Zn transporters, including Slc30a1-10 (also referred to as ZnT1-10) that transport zinc out of the cytosol, and Slc39a1-14 (also referred to as Zip1-14) that transport Zn (as well as Fe and manganese, Mn) into the cytosol(16). Cells also contain metallothioneins (Mt), which serve to bind and buffer Zn ions, and a metal-dependent transcription factor (Mtf1) that translocates to the nucleus upon binding excess Zn to regulate the expression of genes that lower cytosolic Zn levels. One major role of Mtf1 is regulating the expression of Mt genes in response to cytosolic Zn levels, protecting cells from fluctuations in environmental Zn(17). Consequently, increased Mt expression is often used as a proxy for increased cytosolic Zn.

How macrophages use Zn to fight infection seems to depend on the pathogen it encounters, as well as the nature of the host cell. Macrophages poison *M. tb, S. pneumoniae, E. coli, and H. pylori* by transporting toxic levels of Zn into pathogen-containing phagosomes(18–21). Conversely, challenge with *Histoplasma capsulatum* induces macrophages to sequester Zn and increase ROS production to counter the infection(22). There does not appear to be consensus regarding Zn manipulation upon infection with *Salmonella* Typhimurium. *Salmonella* possess genes that aid in surviving Zn toxicity and starvation(23–26), and studies done in different types of macrophage have suggested that Zn can both facilitate and impair infection. In a study of macrophages derived from primary human monocytes and the human monocyte THP-1 cell line, infection induces Zn to accumulate in punctate vesicular compartments, though *Salmonella* managed to avoid these toxic Zn compartments via an SPI-1-dependent mechanism(27). In contrast, *Salmonella* infection of the mouse RAW264.7 macrophage cell line induced Zn mobilization, and increased Zn correlated with impaired bacterial clearance(26). On the other hand, multiple studies have revealed that the high-affinity ZnuABC zinc uptake system and associated accessory proteins are critical virulence factors in *Salmonella*, suggesting that *Salmonella* experience Zn starvation inside the host(23,24,28). Critical comparison of Zn at the host-pathogen interface may be confounded by the differences in the macrophage models used across studies. For example, both Balb/c and C57BL/6 mice lack a functional Slc11a1 (Nramp-1), a phagosomal transporter for magnesium (Mg), Fe, Mn, and Zn(29). Macrophages from these mice are more susceptible to infection and have a reduced capacity to clear intracellular pathogens(30). RAW264.7 cells, a popular mouse macrophage cell line used to study intracellular pathogens, are derived from C57BL/6 mice and therefore equally impaired. The lack of functional Slc11a1 is also likely to compromise the host response to infection with respect to metal regulation, making it difficult to define the nutritional immunity landscape in an immune-competent model system.

In this study, we carried out global RNA sequencing upon infection of macrophages from 129S6 mice with *Salmonella* Typhimurium and examined the changes in gene expression via clustering, and gene set enrichment analysis (GSEA). We deliberately used the 129S6/SvEvTac (hereafter referred to as 129S6) mouse because it contains a functional Slc11a1 protein and has been used as a model system for chronic systemic infection(30). We examined changes in metal-regulatory and metal-dependent genes and compared our results to published data on infection of C57BL/6 mice. While some changes in metal regulatory genes were similar between the two different model systems, suggesting common mechanisms of altering metals at the host-pathogen interface, there were also notable differences, indicating that functional Slc11a1 does alter the nutritional immunity landscape within the host. We also found that cytosolic Zn levels increased over the course of infection and that Zn availability alters infection outcome. Specifically, we found that Zn supports increased replication of intracellular bacteria, but also facilitates clearance of bacteria from macrophages. Our results indicate that functional Slc11a1 significantly affects changes in gene expression, including the immune response, metal regulation in response to infection, and infection outcomes.

## Results

### Infection induces widespread changes in gene expression

To study metal homeostasis during infection, we used bone marrow derived macrophages (BMDMs) from 129S6 mice containing functional Slc11a1. Slc11a1 plays a critical role in regulating metal homeostasis and is important for allowing macrophages to mount an effective immune response to bacterial pathogens(29, 30). To distinguish changes in macrophage gene expression in response to *Salmonella* infection from gene expression changes due to bacterial exposure, macrophages were treated with live or heat-killed *Salmonella enterica* serovar Typhimurium (hereafter referred to as *Salmonella*). Macrophages that were subjected to media changes but not exposed to *Salmonella* served as a control. To examine both early and late immune responses, macrophages were lysed at 2 or 18 hours post bacterial exposure and subjected to global RNA sequencing (Fig 1A). Principal component analysis shows that samples clustered predominantly by time post infection and status of bacteria (alive versus heat killed, Fig 1B).

**Figure 1:**
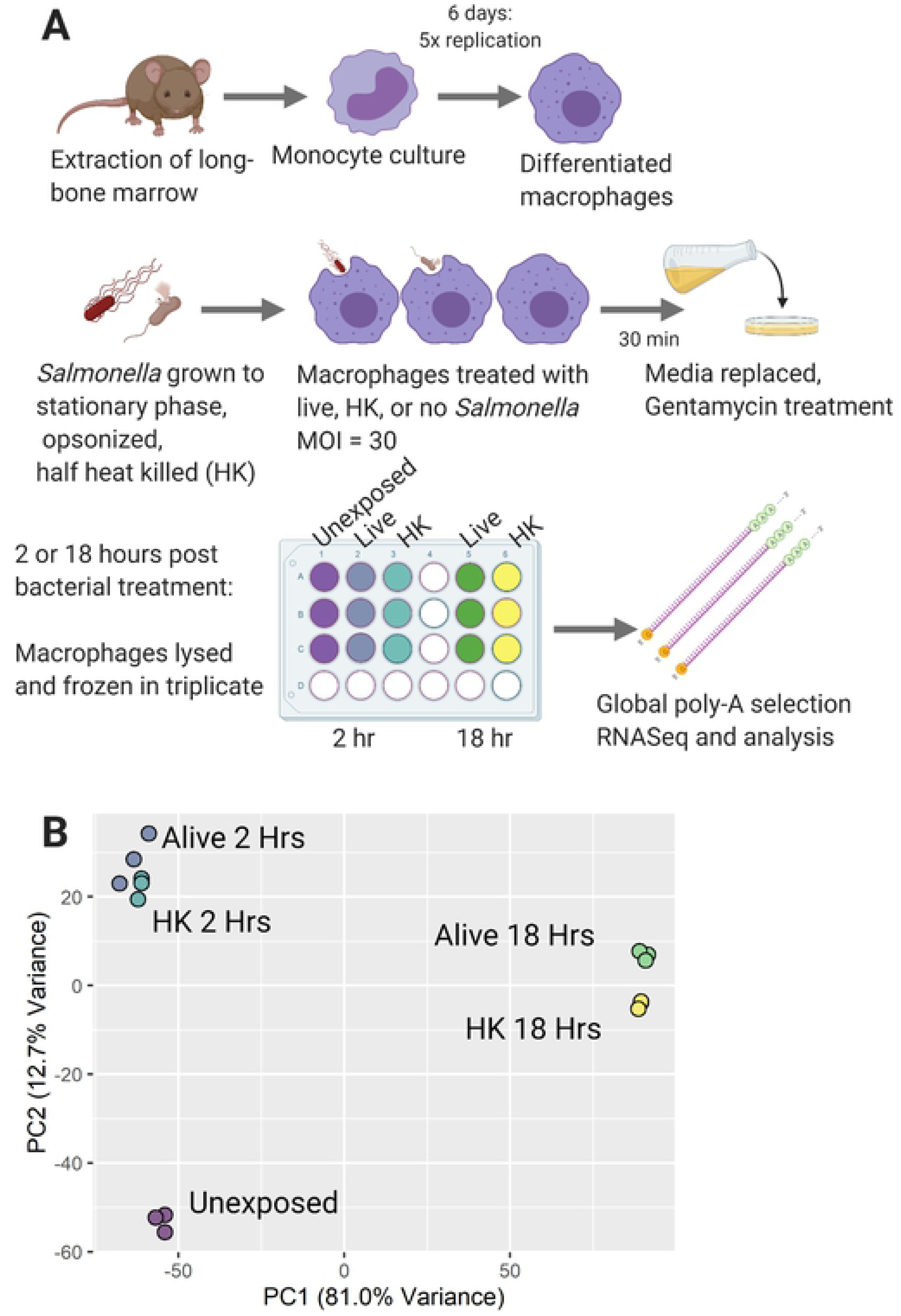
Salmonella treatment of macrophages induces global RNA expression changes across two primary axes: time post exposure and *Salmonella* status (live vs. heat killed) A) Schematic of infection experimental design for 129S6 BMDMs exposed to live or heat killed *Salmonella* Typhimurium SL1344. B) Global principal component analysis (PCA) with x-axis showing differential expression correlated with time post exposure and y-axis showing differential expression correlated with *Salmonella* status (live, heat killed (HK) or unexposed (UE)).

We identified 7766 genes that were differentially expressed and performed hierarchical clustering to examine the main patterns of gene expression changes. We limited analysis to genes with significant differential expression (adjusted p-value (padj) < 0.01), as determined by a DESeq2 log ratio test. An expression level cutoff was also applied (see Methods). An unsupervised algorithm (degPatterns function in DEGreport R package) was used to cluster genes based on similar expression profiles across all conditions. Twenty groups, each with at least 15 genes, emerged. Some groups showed time dependent expression, with little change between bacterial conditions, while others showed significant differences in expression upon treatment with alive versus heat-killed bacteria. Additionally, some groups were different based primarily on expression levels in control cells.

In Fig 2 we highlight the four different expression patterns that encompass the largest gene groups. The remaining groups are presented in **Supplementary Fig S1, S2, S3, and S4** and the genes associated with each group are presented in **Supplementary Dataset 1**. We used DAVID Bioinformatics Resources 6.8 to annotate molecular functions and biological processes associated with the different gene expression clusters. Fig 2A presents group 1, which consists of genes with time-dependent increases in expression. These genes respond to the presence of *Salmonella* regardless of its live or heat-killed (HK) status, and are enriched for mitochondrial function (e.g. oxidative phosphorylation, ATP production), ER-Golgi transport and protein folding, immune response, sugar metabolism, and protein degradation (including proteasome and unfolded protein response genes). Specific genes that are characteristic of the M2 anti-inflammatory immune response(31) such as *il4rα*, *arg1*, *timp1,* and *fcgr2b* are present in this group, along with master regulators of the lipopolysaccharide (LPS) immune response *ifit1*, *stat2*, and *irf7*. Finally, genes involved in pH and redox homeostasis such as *sod1*, *nos2*, *hif1*α, carbonic anhydrases *car2/4/13*, as well as metal regulatory *slc11a1*, iron sequestering *lcn2*, and ferritin iron storage *fth1* are found in this group. Finally, *s100a8*, a metal-binding protein that is a component of calprotectin and plays a prominent role in nutritional immunity exhibited a similar expression pattern, with low expression at 2 Hrs and a significant increase at 18 Hrs (**Supplementary Fig S1D**). These results indicate that the M2 anti-inflammatory response along with many metal associated genes are upregulated over time in response to infection, regardless of whether the bacteria are alive or HK.

**Figure 2:**
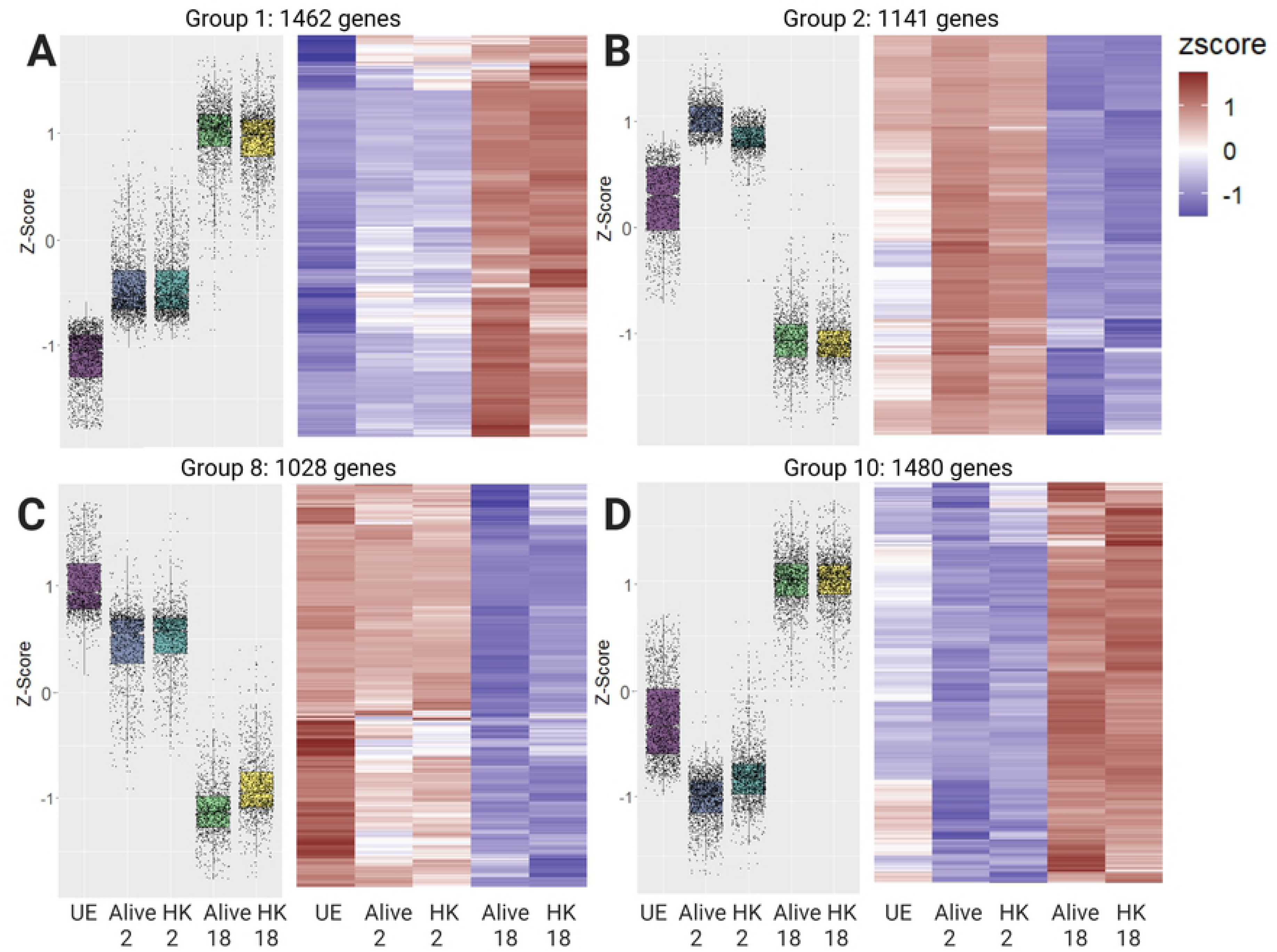
Clustered data exposes multiple distinct expression trends in global data. Four of the twenty groups that emerged from clustering analysis are presented as a box and whisker plot of gene expression levels as Z-scores (normalized across all DE genes) and a hierarchically clustered heat map of gene expression as rlog-scaled DESeq2 normalized counts. A) Group 1 contains 1462 genes that increase in expression at 2 Hrs and increase further at 18 Hrs, without significant differences between the Alive and HK conditions. B) Group 2 contains 1141 genes that increase in expression at 2 Hrs and then decrease below baseline at 18 Hrs. C) Group 8 contains 1028 genes which are expressed in unexposed cells, but then decrease as a function of time post infection. D) Group 10 contains 1480 genes that decrease in expression at 2 Hrs and increase at 18 Hrs. UE: unexposed control cells; HK: stimulation with heat killed bacteria.

The genes in group 2 (Fig 2B) are upregulated at 2 Hrs, with higher expression in alive compared to HK conditions, and are subsequently suppressed below baseline at 18 Hrs. This group is enriched in proinflammatory M1 immune genes(31), including *tnf*, the inflammasome component *nlrp3*, and *irf1* responsible for activating cytokine production. This group also contains multiple genes involved in membrane transport of zinc (*slc30a4*, *slc39a6/a8/a10*). Another hallmark proinflammatory M1 marker *tlr2* and metal regulatory genes (the zinc-dependent transcription factor *mtf1* and iron storage ferritin light chain *ftl1*) are present in group 3 (**Supplementary Fig S1A**, which is similar to group 2 across time but differs in alive vs HK expression at 2 Hrs. Combined, these two groups indicate that in macrophages with functional *slc11a1*, much of the pro-inflammatory response is activated at early time points and then suppressed later in infection.

Fig 2C (group 8) shows genes expressed in the absence of bacterial exposure and distinctly downregulated upon infection over time. This group is notable for its relatively small number of GO annotations, which are focused on cell cycle, cell motility, zinc-binding and transport, and transcription. Noteworthy genes in group 8 include cytosolic zinc exporters *slc30a1/a5* and iron exporter *slc40a1*. Finally, Fig 2D demonstrates group 10 in which genes are suppressed at 2 Hrs post-infection and upregulated above untreated cells at 18 Hrs. This group is enriched for metabolism, protein export, and transcription. Specifically, these genes are involved in ER and Golgi trafficking, respiration, lysosomal activity, zinc transport (*slc39a3/a7*), iron-dependent functions (cytochrome bs, glutaredoxin2, NADH dehydrogenase), mRNA and phospholipid transport, and autophagy. Approximately 10% of these genes are involved in transcription, and half of these transcription factors bind zinc. Also in this cluster are key inflammation-related genes *tlr4* which binds lipopolysaccharide (LPS) and *irf2/3/10* which mediate the Type I IFN response. Combined, these gene expression patterns reveal remodeling of metal homeostasis, metabolism and transcription during infection.

### GSEA reveals pathways differentially affected by live versus heat-killed bacteria

In analyzing global transcription patterns, we found that some groups contain differences in gene expression between live and heat killed bacterial treatments (see groups 4, 5, 7 and 16 as examples), revealing genes that respond differently to a generalized LPS trigger (HK bacteria) vs *Salmonella* with active virulence mechanisms. To identify biological processes or signaling pathways underlying these differences, GSEA(32) was performed on differentially expressed ranked gene list from Alive vs HK at 2 hours, and Alive vs HK at 18 hours. Several significant gene sets were enriched in macrophages treated with live versus HK *Salmonella*, as defined by a false discovery rate of q < 0.05. When using the Hallmark Gene Set from the Molecular Signatures Database(33), we found 47 gene sets enriched at 2 Hrs, and 14 gene sets enriched at 18 Hrs for live versus HK treatment (**Supplementary Dataset S2**). One gene set that is significantly enriched in live *Salmonella* vs HK exposure at both time points is hypoxia (Fig 3A, 3B). Hypoxia is an important aspect of the *in vivo* immune response to infection as hypoxia accompanies inflammation, and a hypoxic environment may affect macrophage M1/M2 polarization(31, 34). Hypoxia is also accompanied by decreased glucose, increased lactate, and decreased pH, and as discussed above, we observed an increase in *hif1*α, carbonic anhydrases, and *nos2*. Multiple studies have shown activation of hypoxia induced factors upon infection of macrophages, and hypoxia is generally associated with macrophage stress as well as defense against bacterial replication(34–36). However, prolonged hypoxia is associated with tissue damage, so the severity and duration of hypoxia is an important factor in infection.

**Figure 3:**
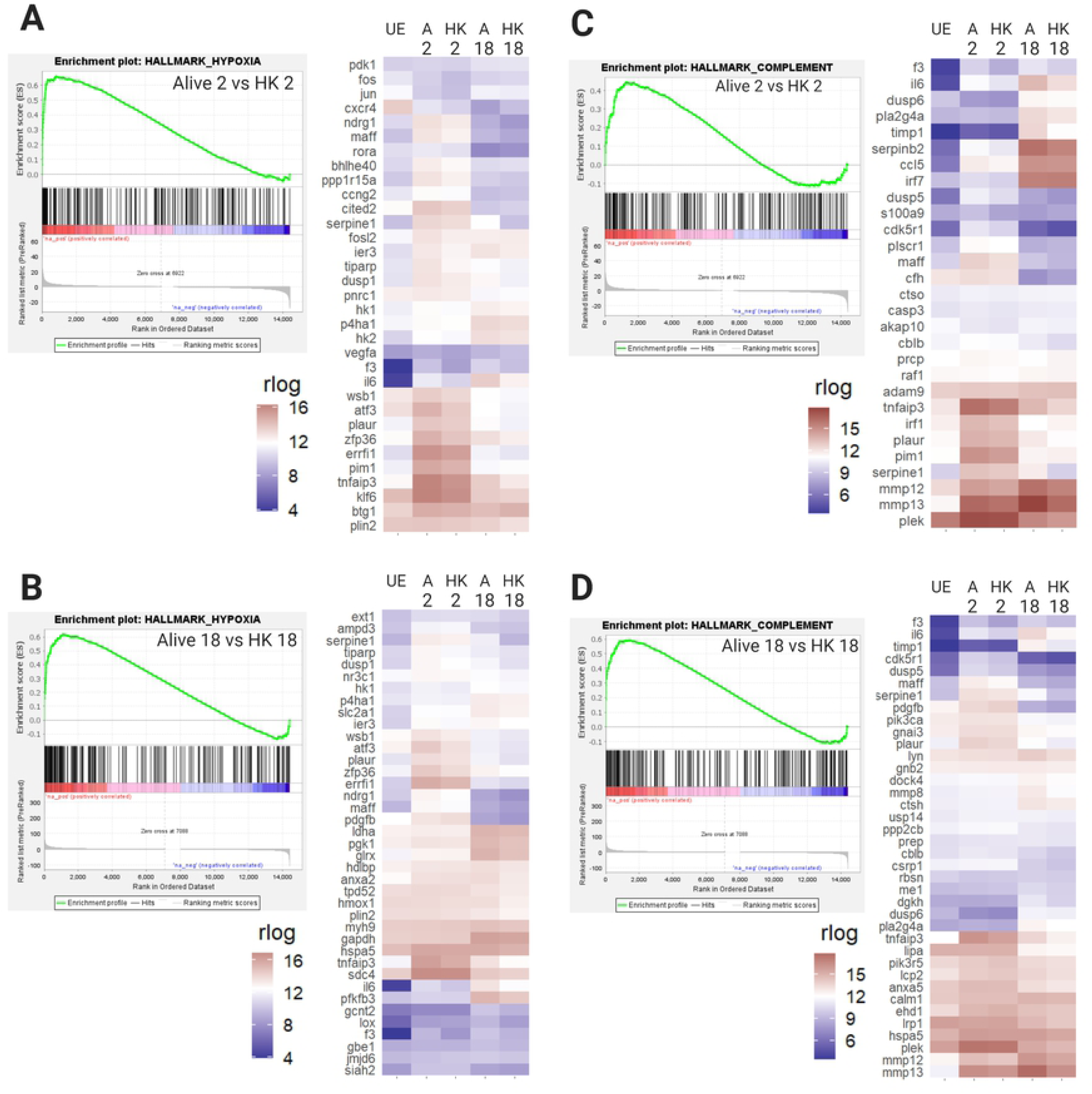
GSEA reveals enrichment of the hypoxia and complement Hallmark gene sets in BMDMs infected with live *Salmonella* compared to treatment with heat killed bacteria. Each panel shows a GSEA enrichment plot for a Hallmark gene set and a hierarchically clustered expression heat map of the leading edge genes from that GSEA plot. In GSEA plots each gene is represented by a vertical black line, and leading edge genes are on the positive end of the spectrum, up to the point of maximum Enrichment Score. **A)** GSEA and heat map of leading edge genes for the Hallmark hypoxia gene set enriched in the 2 Hr Alive condition compared to 2 Hr HK (q = 0.000, NES = 2.12). **B)** GSEA and heat map of leading edge genes for the Hallmark hypoxia gene set enriched in the 18 Hr Alive condition compared to 18 Hr HK. At 18 Hrs, analysis of the hypoxia gene set shows 50 leading edge genes vs. 36 at 2 Hrs. Only 20 of these genes are in the leading edge at both time points. (q = 0.024, NES = 1.65). **C)** GSEA and heat map of leading edge genes for the Hallmark complement gene set enriched in the 2 Hr Alive condition vs. 2 Hr HK (q = 0.062, NES = 1.44). **D)** GSEA and heat map of leading edge genes for the Hallmark complement gene set enriched in the 18 Hr Alive condition vs. 18 Hr HK (q = 0.042, NES = 1.57). 42 complement genes appear in the leading edge, compared to 36 at 2 Hrs. While there is a core set of 16 genes upregulated at both time points, the remaining genes are unique. UE: unexposed control; HK: heat killed; q: false discovery rate; NES: Normalized Enrichment Score from GSEA.

The complement gene set is activated more strongly in macrophages infected with live *Salmonella*. The complement system is part of the innate immune response and mediates a host responses such as opsonization, inflammation and direct bacterial lysis(37). Complement systems respond to bacterial coat components(38) and were therefore expected to respond similarly to live and HK *Salmonella*. Instead, many of these genes showed upregulation in live vs HK conditions (Fig 3C, 3D). At two Hrs post infection, the complement leading edge is enriched for zymogens, enzymes, and secreted proteins, while at 18 Hrs there are more chemokine-and Fc-signaling related genes. At both timepoints, live conditions show upregulation of matrix metalloproteinases *mmp12* and *mmp13* and their inhibitor *timp1*. Additionally, this gene set includes members of the coagulation pathway (*plaur*, *serpin*, *f3*) which overlaps and interacts with complement to trap and kill microbes. Bacteria co-evolved with the complement system, and many bacteria, including *Salmonella*, possess arsenals of anti-complement proteins. Frequently anti-complement proteins work directly against host proteins, but our data suggest live *Salmonella* affect complement gene transcription.

TNFα signaling via NF-κB is also more strongly activated by live *Salmonella*. During infection multiple signaling cascades activate the transcription factor NF-κB, including the proinflammatory cytokine TNFα (TNF in mice). NF-κB is a master regulator and specifically responds to TNF activation with pro-inflammatory and anti-apoptotic signaling. At both 2 Hrs and 18 Hrs, live infected samples were enriched for pro-inflammatory genes, while each time point was enriched in a different set of anti-apoptotic genes **(Supplementary Fig S5)**. At two Hrs *btg2*, *egr3*, *jun*, *vegfa*, *klf4* and *nr4a* were enriched in alive conditions compared to HK, while at 18 Hrs *bcl6*, *smad3*, *stat5a*, *plaur* and *socs3* were upregulated in alive versus HK. Most genes present in the leading edge for both time points were more highly expressed at 2 Hrs, reinforcing the trend of macrophages moving toward an M2-like state later in infection.

We also observed gene sets that were enriched only at 2 Hrs or 18 Hrs. For example, the Hallmark Apoptosis gene set was enriched at 2 hours, and Hallmark Unfolded Protein Response (UPR) was enriched at 18 hours (Fig 4). Late log phase growth induces maximal expression of *Salmonella* Pathogenicity Island 1 (SPI1) genes which allow *Salmonella* to be highly invasive and frequently induce apoptosis in macrophages(39). To promote longer term infections, the *Salmonella* in this experiment were grown to stationary phase before infection, which reduces their expression of SPI1 genes. However, several pro-apoptosis genes are upregulated in Alive exposure at 2 Hrs including the cAMP-dependent transcription factor *atf3*, *jun*, and *pmaip1*. The fact that the apoptotic pathway is not enriched at 18 Hrs suggests that the macrophages manage this response. While UPR and apoptosis are both possible responses to metabolic and ER stress, the upregulation of UPR at 18 Hrs could represent an attempt to restore cellular homeostasis. The transition from apoptosis genes at 2 Hrs to UPR genes at 18 Hrs correlates with a shift in macrophage polarization and may imply decreased ER stress over time.

**Figure 4:**
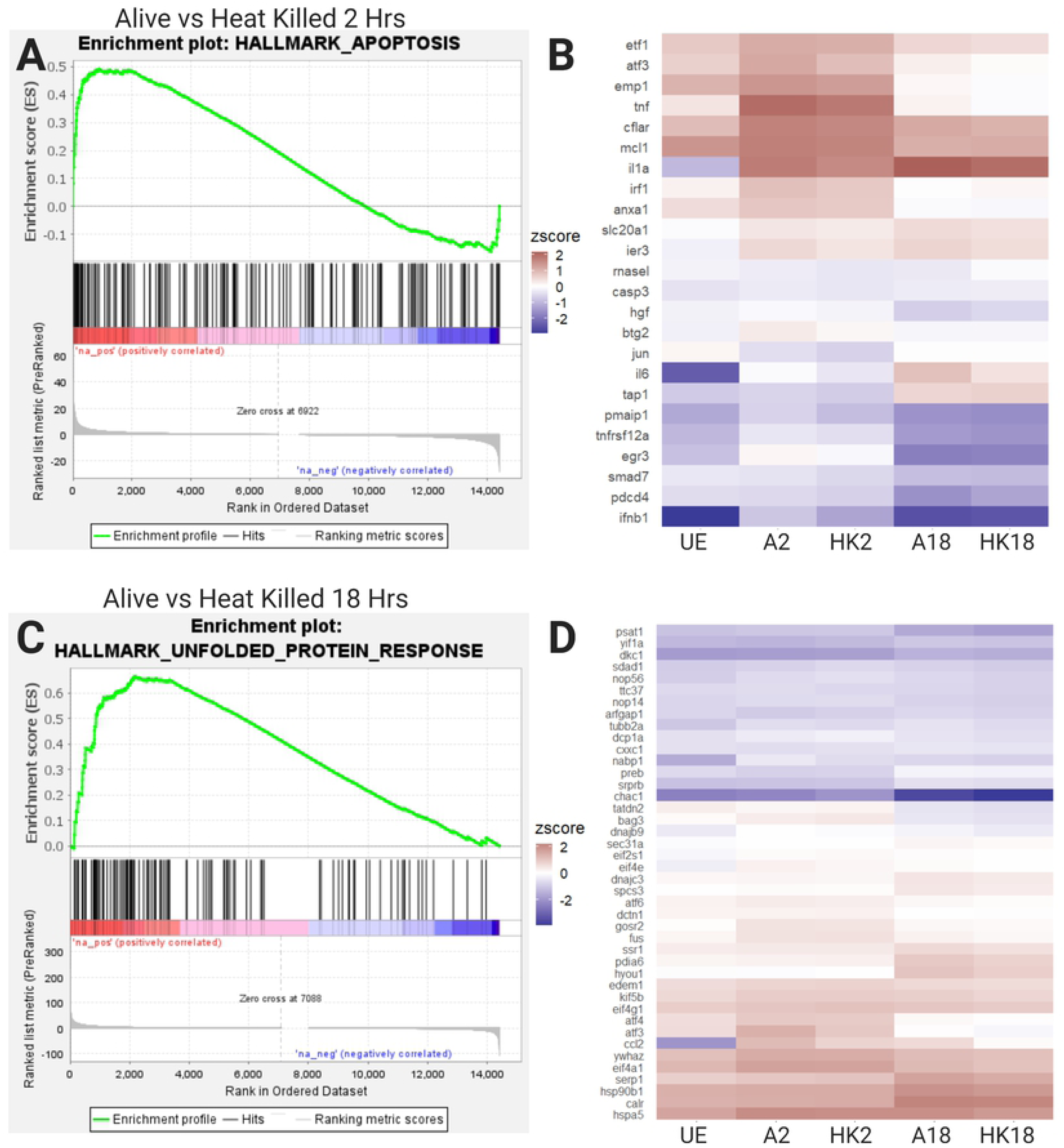
GSEA reveals enrichment of the apoptosis Hallmark gene set at 2 Hrs and the UPR gene set at 18 Hrs in BMDMs treated with live versus heat killed *Salmonella*. **A)** GSEA curve of the Hallmark apoptosis gene set, showing enrichment in the 2 Hr alive condition vs. the 2 Hr HK condition (q = 0.023, NES = 1.58). **B)** Heatmap of 25 leading edge apoptosis genes. **C)** GSEA curve of the Hallmark UPR gene set, showing enrichment in the 18 Hr Alive condition vs. the 18 Hr HK condition (q = 0.021, NES = 1.69). **D)** Heatmap of the 47 leading edge UPR genes. UE: unexposed control; HK: heat killed; q: false discovery rate; NES: Normalized Enrichment Score from GSEA.

### There is significant remodeling of metal homeostasis in response to infection

One of our primary goals in carrying out RNAseq upon infection of 129S6 BMDMs was to define infection-related changes in transition metal regulatory and dependent genes in a model system with functional Slc11a1 (NRAMP1). This allows us to inform our understanding of nutritional immunity in an immune competent model system. Fig 5A shows several major metal regulatory proteins in macrophages. This includes Slc39a1-14 (Zip) transporters that facilitate entry of Zn, as well as Mn and Fe for some Slc39a members, into the cytosol, Slc30a1-10 (Znt) transporters that remove Zn from the cytosol, the Fe exporter Ferroportin (Slc40a1), and Cu transporters Atp7a and Ctr1/2 (Slc31a1/2). Also shown are metallothioneins which store and buffer cytosolic Zn, and the Fe storage protein ferritin (Fth1 – ferritin heavy chain and Ftl1– ferritin light chain), which is protective against Fe-induced oxidative stress in macrophages.

**Figure 5:**
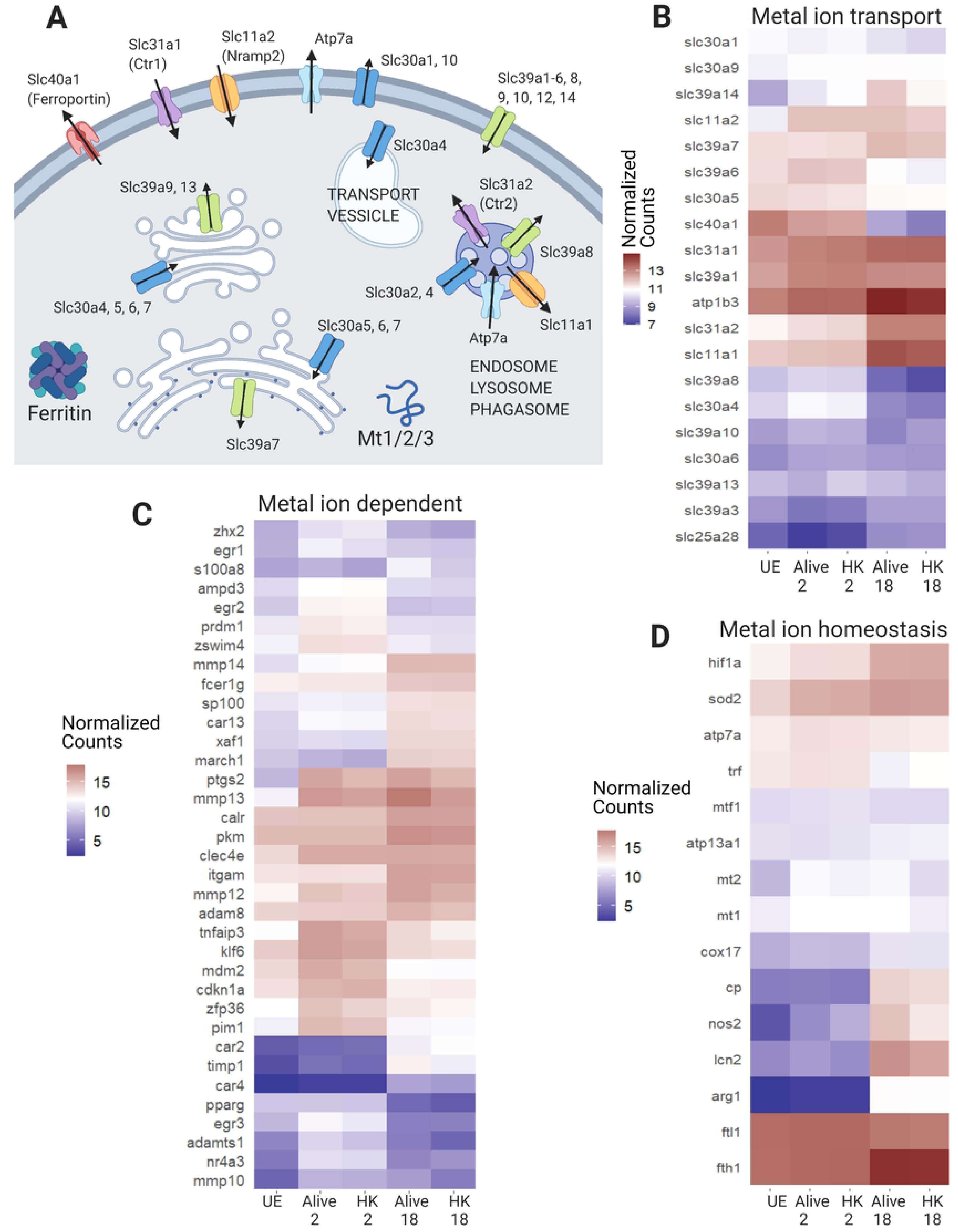
There are widespread changes in metal-transport, -dependent and homeostasis genes. **A)** Schematic of major metal ion transporters and buffering proteins in macrophages. Slc30a1-10 are also referred to as ZnT1-10. Slc39a1-14 are also referred to as Zip1-14. **B-D)** Heatmaps of subsets of metal-transport, -dependent, and -homeostasis genes that are highly expressed and differentially expressed in our data set (log ratio test q < 0.01, mean normalized counts > 100). These gene sets are curated to focus on nutritional immunity metals, including Fe, Zn, Cu, and Mn. Genes that fit in multiple categories are only listed once. **B)** Metal ion transporters. **C)** Metal-dependent and -responsive genes. **D)** Metal ion homeostasis genes. UE: unexposed control cells; HK: heat killed bacteria exposure.

Infection of 129S6 BMDMs leads to broad changes in genes that control metal transport (Fig 5B**, Supplementary Dataset S3**). There is a significant increase in *slc11a1* upon infection, particularly at 18 Hrs. Slc11a1 localizes to the phagosomal membrane where it exports divalent metal cations (Fe, Zn, Mn) while simultaneously acidifying the compartment. The other member of this family, Slc11a2, imports metal ions across the plasma membrane, and is significantly upregulated at 2 Hrs. This may indicate an acute phase emphasis on importing metals into the cell, and a late phase emphasis on exporting metals from the phagosome. At 18 Hrs *slc11a1* is more highly expressed in response to live versus HK *Salmonella*, suggesting that macrophages actively upregulate this gene to help combat live infection.

There are also significant changes in transporters dedicated to import, export, and distribution of Fe, Cu and Zn (Fig 5B). There is a downregulation of the transferrin (*trf*) Fe import pathway and the Fe exporter *slc40a1*, indicating restriction of Fe. The Cu transporters *slc31a1* (plasma membrane) and *slc31a2* (endosome) are significantly increased in expression at 18 Hrs. These changes in gene expression would increase Cu influx into the cytosol, while the upregulation of *atp7a* (endosome) at 2 Hrs suggests an active effort by the macrophages to kill the internalized bacteria. Changes in Zn transporters include upregulation of *slc30a4*, *slc39a14*, *slc39a1*, *slc39a7* and downregulation of *slc30a5*. The significant increase in expression of plasma membrane importer *slc39a14* is especially strong at 18 Hrs, and could result in an increase in cytosolic Zn. Decreased expression of Golgi importer *slc30a5* and increased expression of ER exporter *slc39a7* suggest diminished Zn in the secretory pathway, perhaps to deprive the pathogen of Zn. *Slc30a4*, which transports Zn into the phagosome and secretory vesicles, is upregulated at 2 Hrs and downregulated below control at 18 Hrs, indicating restriction of Zn at late stages of infection. Combined, there are widespread changes in metal transport that indicate limitation of Fe, accumulation of Cu, and redistribution of Zn.

In addition to metal transport, there are significant changes in genes that are metal dependent (Fig 5C) and genes related to metal homeostasis (Fig 5D). There is an increase in intracellular storage capacity for Fe at 18 Hrs; *fth1* increases while *ftl1* drops in expression, reflecting the fact that during infection, heavy chains polymerize together to the exclusion of light chains(40). Similarly, Zn buffering capacity in the form of cytosolic metallothioneins (*mt1/2*) is increased early in infection and maintained with expression at 18 Hrs higher in macrophages treated with live versus HK *Salmonella*. These changes in metallothionein would be consistent with increased cytosolic Zn late in infection. There is also a significant increase in expression of Zn-dependent carbonic anhydrases that regulate pH, Zn-dependent proteases (*mmp10*, *12, 13, 14, timp1, adamts1, adam8*), the ferroxidase ceruloplasmin (Fig 5C). Additionally, there is an increase in expression of lipocalin, a protein involved in innate immunity by binding Fe-bound siderophores and depriving pathogens of Fe. Finally, there are significant increases in genes involved in metal homeostasis and oxidative stress, particularly at late stages of infection. These include: *hif1a*, Cu/Zn-dependent *sod2* and *sod3*, Mn-dependent *arg1*, and Zn-dependent *nos2*. These changes indicate that remodeling metal transport is accompanied by significant changes in the expression of metal-dependent enzymes and metal storage.

### Cytosolic labile Zn increases at late stages of infection

The widespread changes in Zn transporter expression upon infection suggest remodeling of Zn homeostasis, and would be expected to lead to an increase in cytosolic Zn and decreased Zn in the secretory pathway. Previous studies in different model systems have suggested different changes in intracellular labile Zn pools in response to infection(27, 41). However, all previous studies have used small molecule indicators such as FluoZin3 or Zinpyr-1, which often exhibit variable and unpredictable localization within a cell and can confound measurement of Zn in a defined location. For example, FluoZin3-AM has been shown to localize to the cytosol, Golgi, lysosomes and vesicles, and localizations differ among cell types(10,42,43). Previous studies in Slc11a1 defective Raw264.7 cells used FluoZin3 to determine that there was an increase in labile Zn upon infection via FACS. Unfortunately, the absence of fluorescence images precluded evaluation of where the Zn increase occurred. In human monocyte-derived macrophages FluoZin3 appears to be in vesicles or early endosomes(27), and reported an increase in Zn in these compartments upon infection. Of the studies that have attempted to measure Zn inside activated or infected macrophage, none have unambiguously addressed whether there are changes in cytosolic Zn.

To directly measure cytosolic Zn in 129S6 BMDMs at various time points post-infection, we used the genetically encoded zinc sensor NES-ZapCV2(44, 45). This sensor uses the ratio of YFP emission to CFP emission upon CFP excitation (Forster Resonance Energy Transfer, FRET, ratio) to measure labile Zn in cells. The advantage of the genetically encoded sensor is that localization of this sensor is restricted to the cytosol. Briefly, cells were nucleofected with a plasmid encoding NES-ZapCV2 15 hours prior to infection with *Salmonella*. At select time points post-infection cells were imaged on a widefield fluorescence microscope to measure the resting FRET ratio (Fig 6A). Cells were subsequently subjected to an *in situ* calibration to determine the minimum and maximum FRET ratio in each cell (Fig 6B) and the calibration parameters were used to calculate the fractional saturation of each cell under resting conditions and the concentration of labile cytosolic Zn (see Methods for details, Fig 6C). Uninfected cells have a mean fractional saturation of 0.30 which corresponds to a Zn concentration of 80 pM. At 12 Hrs and beyond, the fractional saturation increased to ∼ 0.5 which corresponds to ∼ 2.9 nM Zn. Our data show that labile Zn in the cytosol does indeed increase later in the infection (36-fold), consistent with the observed increase in expression of *mt1* and *mt2*.

**Figure 6:**
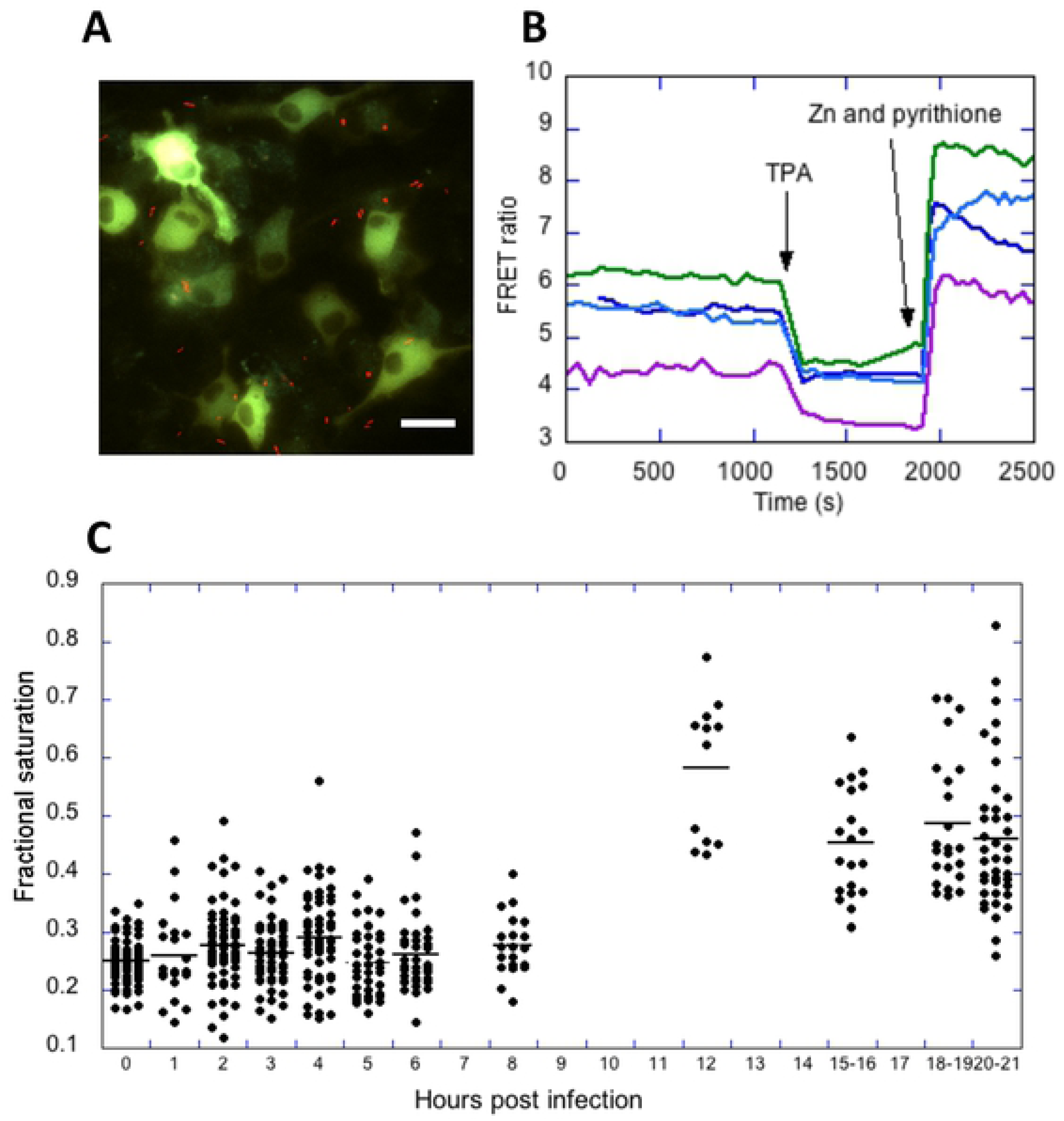
Changes in labile cytosolic Zn in 129S6 BMDMs upon infection with *Salmonella*. **A)** Representative image of BMDMs expressing the NES-ZapCV2 fluorescent Zn sensor (green) treated with *Salmonella* constitutively expressing mCherry (red). Scale bar = 20 μm. B) Representative *in situ* calibration of the NES-ZapCV2 sensor showing the FRET ratio (R, background corrected FRET channel divided by the background corrected CFP channel) over time. The R_min_ is obtained upon addition of 50µM TPA and the R_max_ is obtained upon addition of 23.8nM buffered Zn, 0.001% saponin and 0.75µM pyrithione. Each trace represents a single cell. C) Fractional saturation of the NES-ZapCV2 sensor at different time points post infection. Each dot represents an individual cell. Data represent at least two independent experiments per time point.

### Zn promotes bacterial clearance as well as bacterial replication

Having established that Zn regulation and Zn availability in the cytosol are altered upon infection, we evaluated the influence of Zn availability on infection outcome. While a previous study examined increases and decreases in Zn on *Salmonella* infection of Raw264.7 cells(26), we explore the impact of Zn on infection outcome in immune competent macrophages with functional Slc11a1. To examine infection outcomes, we incorporated the pDiGc plasmid (Fig 7A) into *Salmonella*, which encodes two fluorescent proteins. DsRed is expressed under an arabinose inducible promoter while GFP is constitutively expressed(46). This plasmid allows tracking of infected cells (green signal) and bacterial replication (dilution of the red signal upon cell division). Arabinose induction of DsRed is halted upon macrophage infection, such that any bacterial division occurring within the macrophage leads to dilution of the red fluorescent signal. Comparison of green and red signals across time yields a picture of bacterial division vs macrophage clearance of bacteria.

**Figure 7:**
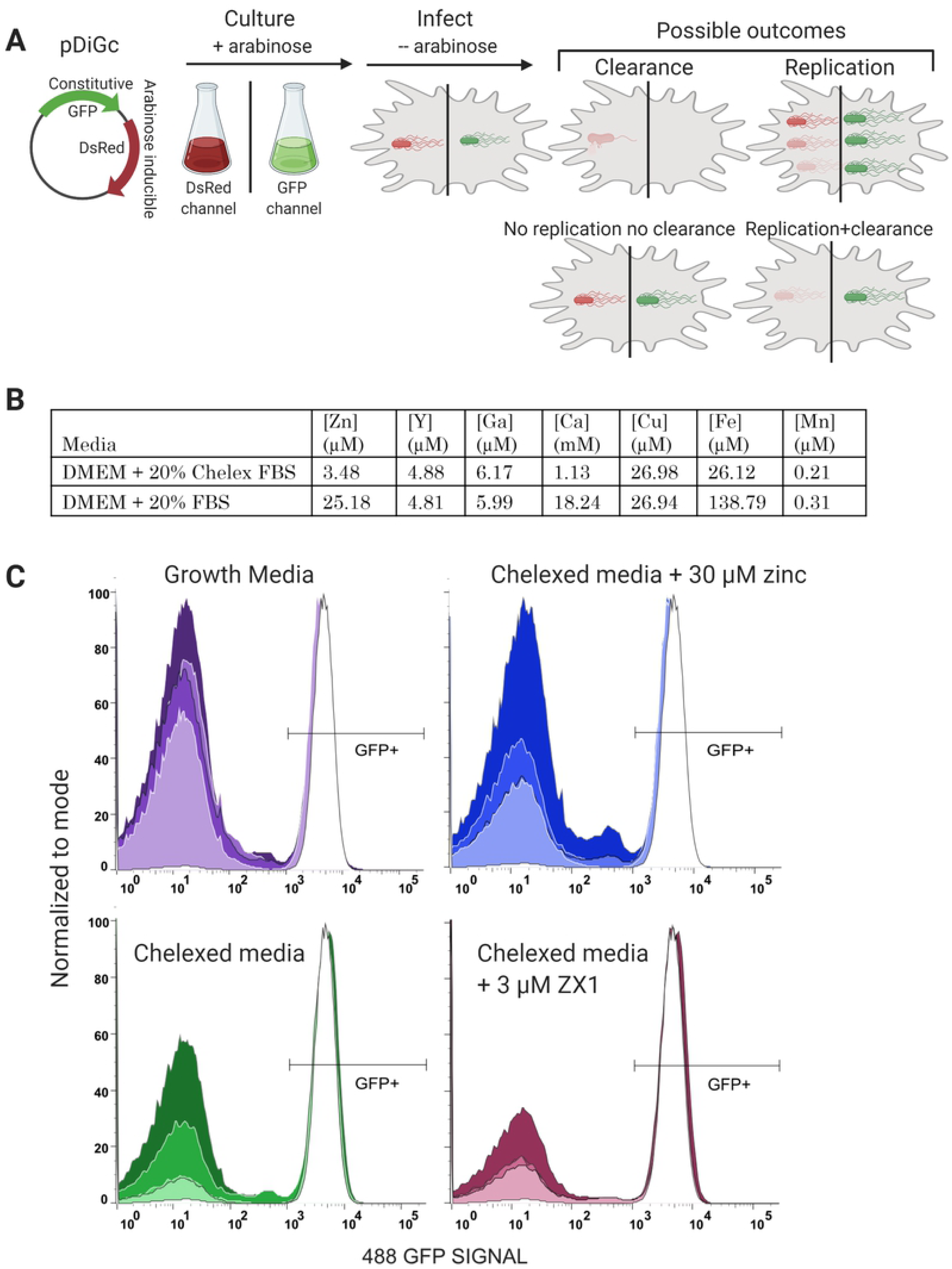
Bacterial clearance as a function of time and Zn availability. **A)** Schematic of the experimental protocol. Salmonella expressing the pDiGc plasmid expresses GFP constitutively and DSRed under arabinose induction. Bacteria are cultured with arabinose and aeration to the stationary phase, at which point they express both fluorescent proteins. Arabinose is removed at infection, and infected macrophages are fluorescent in both red and green channels by flow cytometry. As infection proceeds the green to red fluorescent ratio ‘tracks’ the bacteria. **B)** Metal content of normal growth media and media containing Chelex-treated FBS, as determined by inductively coupled plasma mass spectrometry (ICP-MS). **C)** Histograms of the GFP signal in macrophages expressing pDiGc in 4 different medias and at 4 time points. Each experimental condition was done in triplicate, and 10,000 cells were measured per sample. Replicates were essentially identical, so data for the replicates was concatenated. Increasing color saturation indicates time post *Salmonella* exposure (2, 10, 18 or 24 hours). The right peak in each graph corresponds to the GFP+ macrophages, indicating infection. As *Salmonella* are cleared the macrophages become GFP-and shift to the left peak on each graph. At the initial inoculation (white curve in each panel), macrophages show almost 100% infection rate. Normal macrophage growth media and Chelexed media supplemented with 30 μM Zn (replete Zn conditions) show almost 100% bacterial clearance at 24 hours. Chelexed media (low Zn) shows decreased bacterial clearance while Chelexed media with 3 μM extracellular zinc chelator ZX1, (essentially no Zn) shows poor bacterial clearance.

To alter Zn availability, we manipulated Zn levels in the macrophage media after *Salmonella* had been internalized. Four media conditions were used in this study to span the range from Zn replete to Zn deficient. The four conditions were: standard macrophage growth media (DMEM + 20% FBS) containing 25 μM Zn (Fig 7B), or media prepared with Chelex-100-treated FBS to deplete metal ions (4 μM Zn, Fig 7B), which was then further depleted of Zn (Chelex media + 3 μM ZX1, ∼ 0 μM Zn) or reconstituted with replete Zn (Chelex media + 30 μM Zn, 34 μM Zn total).

Macrophages were infected with *Salmonella* expressing pDiGc in macrophage growth media and infection proceeded for 45 minutes before experimental media conditions were introduced. This avoided the confounding variable of Zn’s effect on the phagocytosis of bacteria. Infected macrophages in each media condition were fixed at four time points: 2, 10, 18 and 24 Hrs post inoculation and subjected to flow cytometry. A control sample was fixed immediately after the 45 min infection and verifies that 91% of macrophages contained GFP-positive bacteria at the beginning of the experiment. Fig 7C demonstrates the loss of GFP signal in macrophages in each growth media as a function of time. The increase in the fraction of cells with no GFP is indicative of bacterial clearance. In each panel, the white curve is from the initial infection, and increased color saturation indicates time post-infection. In normal growth media, clearance of bacteria is strong at 2 Hrs, with 70% of initially infected cells having lost all GFP signal. Clearance further increases over time, with 76%, 74%, and 85% clearance at 10, 18 and 24 Hrs. Control and Zn replete media contained similar Zn levels but different amounts of Ca and Fe due to Chelex treatment. At early time points (2, 10 and 18 Hrs) there is 56%, 64% and 59% bacterial clearance in the Zn replete media compared to the control, likely due to either depleted Ca or Fe. By 24 Hrs, Zn replete samples reached a similar level of infection clearance, with 87% clearance. In contrast, macrophages in Chelexed media with low Zn and ZX1-treated media with effectively no Zn show significantly impaired bacterial clearance. At 24 Hrs Chelex and Zx1 conditions had achieved 73% and 53% clearance, respectively. Direct comparison of the different media conditions at each time point are presented in **Supplementary Figure S6**. These results confirm that 129S6 macrophages are competent at clearing *Salmonella* throughout infection and that Zn availability positively correlates with the effectiveness of bacterial clearance.

In addition to measuring bacterial clearance, the pDiGc plasmid can also be used to measure bacterial replication, as the DsRed signal is diluted with every cell division. We compared the DsRed signal versus GFP signal as a function of time and different media conditions (Fig 8). In standard macrophage growth media (top row), the DsRed signal decreases over time in 23% of the GFP+ cells, indicative of bacterial replication. It is important to note that although 129S6 macrophage are efficient at clearing the *Salmonella* infection, here we are looking at the small portion of the macrophage population that remains infected (GFP^+^). The Zn replete media (2^nd^ row) shows bacterial replication comparable to standard growth media, with 23% of GFP^+^ macrophages showing diluted DSRed signal at 24 Hrs. In contrast, in Chelexed media with depleted Zn and Chelexed media containing ZX1 (no Zn) there is weaker bacterial replication (11% and 7%, respectively), as indicated by the lack of a fluorescent “tail” at lower DsRed intensity. Interestingly, this result suggests that while Zn facilitates host clearance of bacteria, it also helps bacteria to replicate in cells that do not clear the infection. This may suggest that bacterial clearance occurs more easily in dividing bacteria as persistent infections are mainly non-replicating.

**Figure 8:**
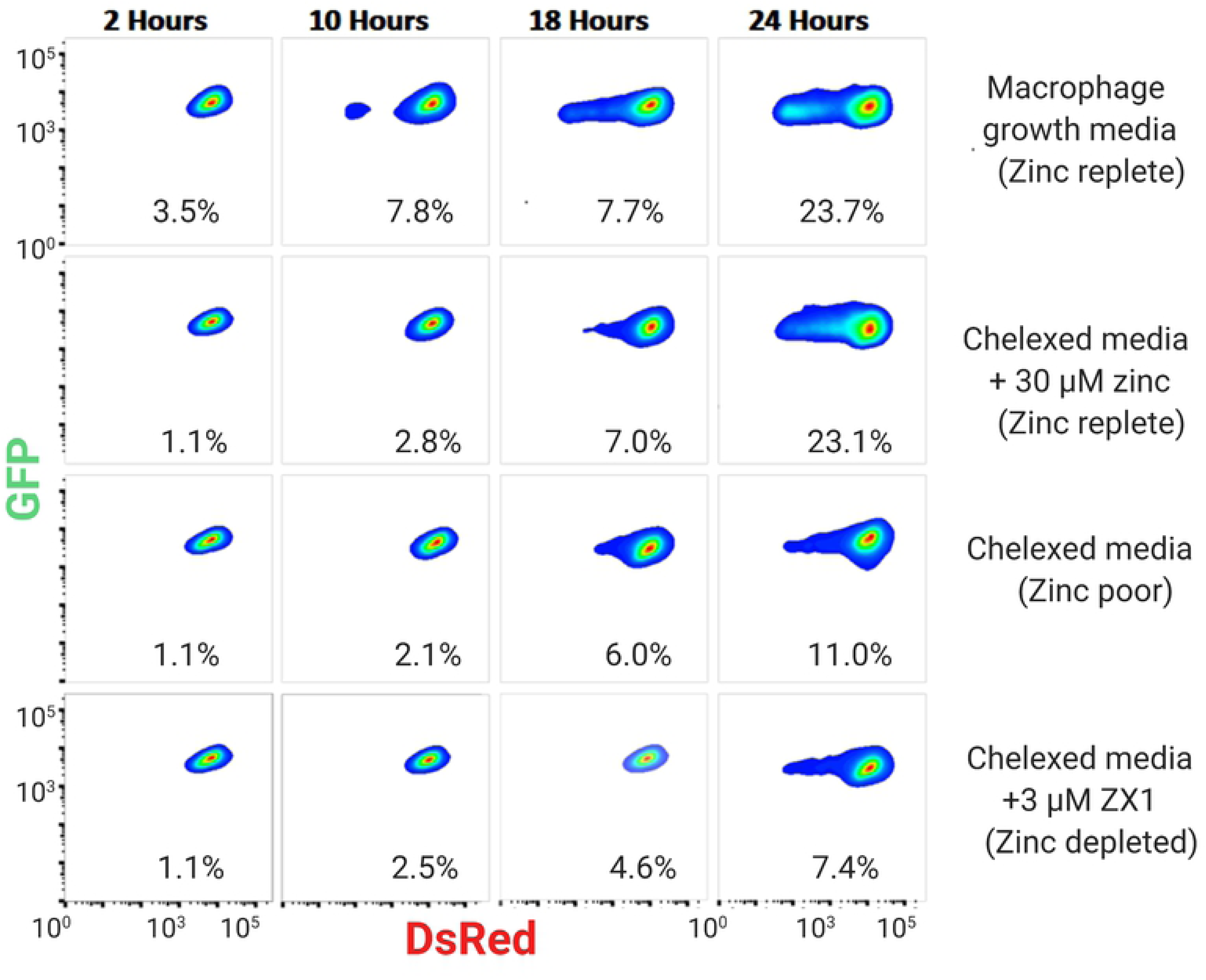
Bacterial replication as a function of time and Zn availability. 129S6 BMDMs were infected with *Salmonella* expressing pDiGc. The GFP and DsRed signals in BMDMs were measured by flow cytometry and plotted against one another. These density graphs show only macrophages containing GFP^+^ *Salmonella*, which indicates active infection. Percentages on each plot indicate the number of macrophages with diluted DSRed signal, an indicator of *Salmonella* division. Dilution of DsRed is a function of time and zinc availability, with all media conditions showing an increase in bacterial division over time, but control and zinc replete conditions (first and second rows) show much more robust division.

### Macrophages from 129S6 and C57BL/6 differ in nutritional immune response to infection

Because the Slc11a1 transporter specifically aids in depriving pathogens of metal ions, we hypothesized that expression of genes related to metal handling in our cells would differ significantly from macrophages derived from Slc11a1 non-functional mice (BALB/c, C57BL/6, and the Raw264.7 cell line). Therefore, we performed a comprehensive analysis of changes in expression in 129S6 macrophages (this study) and macrophages derived from C57BL/6 mice (RNAseq study by Stapels et al(47)) at 18 hours post infection. Specifically, we compared our unexposed control (UE), 18 Hr Alive and 18 Hr HK conditions against uninfected macrophages (UI), those containing growing *Salmonella* (G), and bystander macrophages not containing *Salmonella* at 18 Hrs (BY) from Staples et al. All data were analyzed on the same pipeline and normalized together within DESeq2 to make the data as comparable as possible **(Supplementary Dataset S4)**.

To evaluate potential differences in genes involved in nutritional immunity between the two datasets, we examined all genes with UNIPROT keywords for iron, zinc, or copper transport or homeostasis. Fig 9 presents heat maps for Fe, Cu, Zn transport and Ze-dependent genes. There are a number of similarities in regulation of metal-transport, -homeostasis, and -dependent genes, indicating that there are common mechanisms for managing metal ions at the host-pathogen interface, even when there are differences in the host. In particular, there is a significant decrease in Fe import (*trf*), export (*slc40a1*), and an increase in ferritin storage (*fth1*) in both studies (Fig 9A). Similarly, both studies show upregulation of plasma membrane copper transporter *slc31a1* indicating Cu uptake from the extracellular space (Fig 9B). With respect to Zn regulation, both studies show upregulation of plasma membrane Zn importer *slc39a14*, downregulation of the Golgi importer *slc30a5*, and an increase in expression of Zn buffers *mt1* and *mt2* (Fig 9B, 9C). Combined these results suggest a general pattern of limiting the availability of Fe, increasing exposure to Cu, and increasing cytosolic Zn.

**Figure 9:**
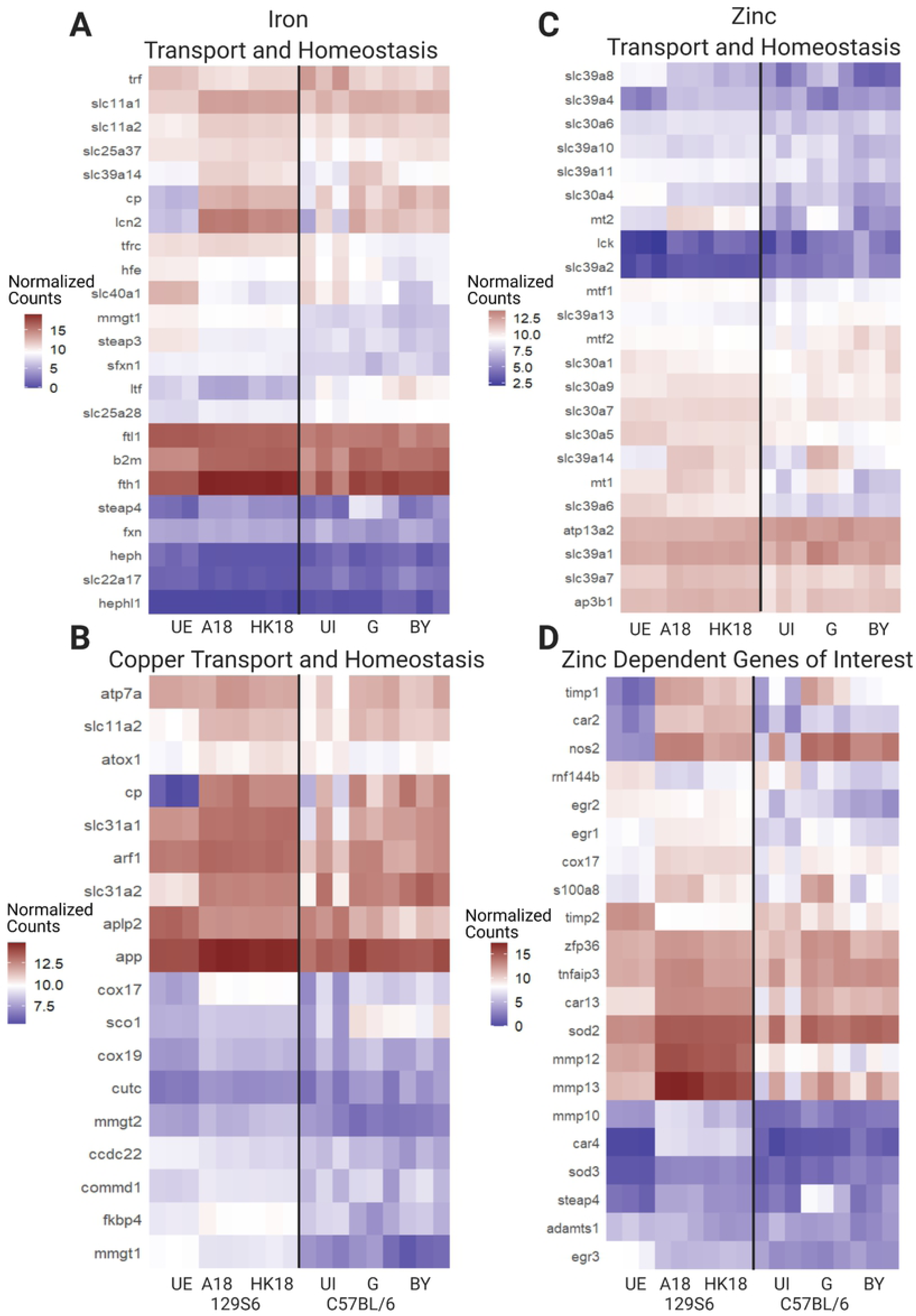
Changes in metal-transport and -homeostasis genes reveal differences in metal regulation in BMDMs derived from Slc11a1-functional mice compared to nonfunctional mice. RNAseq data from this study are compared to data from Stapels et al, `GEO Accession GSE104785`. Data from both studies were processed through the same pipeline and normalized together. Gene lists were compiled using Uniprot keywords and the mouse genome, and filtered for mean normalized count > 5 in at least one data set. A) Genes involved in Fe transport and homeostasis. B) Genes involved in Cu transport and homeostasis. C) Genes involved in Zn transport and homeostasis. UE: 129S6 BMDMs unexposed to *Salmonella*; HK: heat killed *Salmonella* treatment; UI: uninfected C57BL/6 BMDMs; G: C57BL/6 BMDMs containing growing *Salmonella*; BY: bystander C57BL/6 BMDMs.

Despite these similarities, there are also notable differences that suggest the presence of functional *slc11a1* alters metal regulation at the host-pathogen interface. With respect to Fe, lactotransferrin (*ltf*), a major Fe binding factor with bactericidal properties, *steap3*, which encodes a ferrireductase that reduces ferric Fe released from transferrin in the endosome, and *mmgt1*, which encodes a Mg transporter that ferries multiple metal ions across membranes are all downregulated in infected 129S6, but do not respond to infection in C57BL/6 cells. There are also notable differences in key Cu regulatory genes. The Cu importer *slc31a2* is upregulated in 129S6, while there is no change in C57BL/6. Copper chaperones play an important role in distributing Cu and loading it into enzymes; the copper chaperone *atox1* is upregulated in 129S6, but downregulated in C57BL/6; *cox17* is upregulated in 129S6, but not C57BL/6 (q > 0.06). Finally, *sco1* is upregulated in C57BL/6 but not 129S6. There are also differences in the magnitude of gene expression changes, with more marked changes in 129S6, including for ferroportin (*Slc40a1*), multi-copper enzyme ceruloplasmin (*cp*), and *lcn2*, which reduces bacterial replication by sequestering Fe bound to microbial siderophores. *Lcn2* is also intimately involved in regulating the expression of pro-apoptotic *bcl2l11,* which is strongly downregulated in 129S6 but is unaffected in C57BL/6 cells, suggesting these two cell types have disparate apoptotic tendencies at 18 Hrs post infection.

There are also notable differences in infection-induced expression changes in genes encoding Zn transporters and Zn-dependent proteins between the two systems (Fig 9C, D). Specifically, upregulation of *slc39a7* and downregulation of *slc30a4* in 129S6 but not C57BL/6 suggest limitation of Zn in the secretory pathway and phago-lysosomal compartment in response to infection in 129S6 BMDMs. Additionally, *slc39a8* is downregulated in 129S6 but upregulated in C57BL/6. In primary human lung macrophages, the Zn importer *slc39a8* is upregulated in response to LPS and negatively regulates proinflammatory responses(48). Zn-dependent proteins such as proteases (*mmp9/10/12/13*) and carbonic anhydrases (*car 4/11*), are upregulated in 129S6 but not C57BL/6. Finally, there are notable differences in the magnitude of gene expression changes, including *car2*, *car4*, and *nos2*. These differences suggest changes in Zn distribution between the two model systems and altered regulation of Zn-dependent enzymes.

A comparison between A18 vs UE and G vs UI revealed global expression differences as well. 1991 genes are differentially expressed (padj < 0.01) between G and UI that were not differential (padj > 0.03) between A18 and UE. Further, 2723 genes were significantly different between A18 and UE, but not in G versus UI. When these gene lists were scrutinized using DAVID annotation clustering, the genes differential in 129S6 were enriched for cell cycle functions, and the majority are down regulated. Interestingly, the genes differentially expressed in C57BL/6 were enriched for apoptosis, and almost all of these genes were upregulated. These data suggest that these two cell types are moving toward different fates by 18 Hrs post *Salmonella* exposure, with 129S6 cells entering quiescence but C57BL/6 cells experiencing apoptosis. This observation is consistent with the fact that Slc11a1 knockout macrophages are models for acute infection and are more susceptible to killing by *Salmonella*.

## Discussion

This work presents a systematic analysis of changes in global gene expression, as well as a more targeted analysis of changes in genes related to transition metal ions in 129S6 macrophages infected with *Salmonella*. The 129S6 model system was chosen because these mice contain functional Slc11a1 (Nramp1), a divalent metal ion transporter that localizes to phagosomes and regulates metal availability to the intracellular pathogen. Slc11a1 confers resistance to intracellular pathogens and the 129S6 mouse has been used to model long-term chronic infection by *Salmonella*, as the mice don’t succumb to initial acute infection the way C57BL/6 or BALB/c mice do(30,49,50). A recent study examined the *Salmonella* proteome upon infection of mice homozygous or heterozygous for functional versus nonfunctional Scl11a1 and found an abundance of Mg, Zn, Fe and Mn uptake systems expressed in *Salmonella* in resistant mice but not *Salmonella* that colonized susceptible BALB/c mice(49). This study indicates that Slc11a1 plays a major role in affecting metal ion availability for uptake by *Salmonella*. However, how metal regulation in the host differs in the presence and absence of functional Slc11a1 has not been explored. Given that metal manipulation is a key component of nutritional immunity, we sought to define how genes encoding metal transporters, metal-dependent proteins, and metal-regulatory proteins were affected by treatment with live or heat killed *Salmonella* at early (2 Hrs post infection) and late (18 Hrs post infection) time points.

We observed widespread changes in metal transport, metal-dependent, and metal homeostasis genes, suggesting remodeling of Fe, Cu, and Zn availability by host cells. There is significant upregulation of *slc11a1*, especially at late stages of infection, consistent with the host cell limiting Fe, Mg, Mn and Zn in the phagosome. Further changes in gene expression are consistent with decreased Fe uptake (*trf*), export (*slc40a1*) and increased storage (*fth1*); increased Cu uptake (*slc31a1*), import into the phagosome (*atp7a*) and altered distribution (Cu chaperones *atox1, sco1, cox17*) and increased transport of Zn into the cytosol (*slc39a1*, *slc39a14*) coupled with decreased transport into the secretory pathway and endo-lysosomal system (*slc39a7, slc30a4, slc30a5*). Increased expression of *mt1* and *mt2* genes, which encode Zn buffering proteins, further suggests an increase in cytosolic Zn. We also observed numerous changes in metal-dependent proteins and enzymes, many of which play an important role in nutritional immunity (*lcn2, ltf, cp, steap3, sod2, arg1, nos2, mmp, car, s100a8*). Some of these changes have been seen in a variety of mouse model systems in response to diverse intracellular pathogens, suggesting widespread or universal strategies for the host to combat infection. In particular, the changes in gene expression to limit Fe(6, 51) and accumulate Cu appear to be common strategies for nutritional immunity(4). Indeed, the changes in expression of *trf, slc40a1, fth1, slc31a1, lcn2* and *cp* were also observed in analysis of RNAseq data from infection of macrophages from C57BL/6 mice at 18Hrs post infection. However, it is notable that many of the changes are significantly more pronounced (larger fold change) in the 129S6 model system, indicating a more robust nutritional immunity response in macrophages with functional Slc11a1. This observation is consistent with the findings of Cunrath and Bumann, which found that only in infection of mice with functional Slc11a1 were bacterial metal uptake systems induced(49), suggesting that functional Slc11a1 is necessary to activate nutritional immunity mechanisms. Interestingly, the changes we observed in expression in *slc11a1*, as well as key Fe and Cu regulatory genes were comparable in macrophages infected with live bacteria or treated with HK bacteria. This suggests that the host response is driven not only by active pathogen virulence factors, but also pathogen-associated cues, such as LPS.

As noted previously, there does not appear to be a universal strategy for host manipulation of Zn at the host-pathogen interface, with examples of host mechanisms to both starve and poison an intracellular pathogen(2,5,22). Cellular regulation of Zn is more complex than Fe or Cu, with 24 Zn transporters that control transport into and out of the cytosol as well as intracellular organelles. Indeed, pathogens and immune system modulators have been shown to have different effects on Zn transporter expression(7) including upregulation of *slc39a8*(48) in response to LPS and TNFα, upregulation of *slc30a4* and *slc30a7* upon stimulation with GM-CSF(52), and upregulation of *slc39a14* in response to LPS(53). Previous studies have suggested that *Salmonella* experiences Zn starvation inside the host as the high affinity bacterial Zn uptake system ZnuABC(23) and associated accessory protein ZinT(24) and low affinity uptake system ZupT(54) contribute to virulence in mice. But the concept of Zn starvation is hard to reconcile with reports of Zn accumulation in puncta(27) and the suggestion that *Salmonella* induce Zn elevation to evade bacterial clearance(26). To provide a more complete picture of Zn modulation within the host, we analyzed changes in the expression of Zn regulatory genes, measured labile Zn levels in the cytosol at different times post infection, and examined the effect of Zn availability (based on manipulation of Zn levels in media) on infection outcomes: bacterial clearance and bacterial replication. We were particularly interested in comparing how the host cell remodels Zn availability in macrophages with functional Slc11a1, and hence a robust nutritional immunity response.

The changes in Zn transporters and homeostasis genes predict an increase in cytosolic Zn and a decrease in Zn in the ER / Golgi / phagosome. Specifically, we observed significant upregulation of *slc39a1*, *slc39a14*, *mt1* and *mt2*, with more pronounced changes at late stages of infection (18 Hrs). Slc39a1 and a14 are plasma membrane transporters that mediate entry of Zn into the cytosol, while Mt1 and Mt2 are Zn buffering proteins that are frequently used as proxies for cytosolic Zn, as increased Zn leads to increased *mt1/2* expression via the Zn-dependent transcription factor Mtf1. Using a genetically encoded Zn sensor explicitly targeted to the cytosol, we measured an increase in Zn from ∼ 80 pM to ∼ 2 nM at 12 Hrs post infection that remained elevated through the last measurement time (21 Hrs post infection). Intriguingly, *mt1*, *mt2*, and *slc39a14* are more highly upregulated upon infection with live *Salmonella* compared with heat killed bacteria, suggesting an active recognition of and response to live bacteria. We speculate that the host cell increases cytosolic Zn to support metabolic and transcriptional proteins that fight infection. This is consistent with our observation that a large fraction of differentially expressed genes has DAVID annotations that include “Zn binding”. When we manipulated Zn availability in the media, we found that Zn replete conditions led to robust clearance of infected bacteria, suggesting that the host may use elevated cytosolic Zn to fight infection.

We also observed an increase in *slc39a7* which encodes the transporter that moves Zn out of the ER and secretory pathway, a decrease in *slc30a5* which encodes the transporter that moves Zn into the Golgi and vesicular compartments, and a decrease in *slc30a4* which encodes a transporter which moves Zn into the phagosome. All three of these changes were comparable in macrophages treated with live *Salmonella* versus heat HK bacteria, suggesting a general host response to the pathogen and pathogen-associated cues (for example LPS). Further, all three of these changes in expression would lead to Zn limitation in the secretory / phago/lyoso-somal pathway and hence Zn restriction for the pathogen. This finding is consistent with previous observations that *Salmonella* upregulate bacterial Zn uptake mechanisms, especially in the presence of functional Slc11a1. Manipulation of Zn in the media revealed that zinc availability strongly correlated with bacterial replication, where Zn limitation blocked replication and Zn elevation enabled replication. Combined, our results indicate that manipulation of Zn at the host-pathogen interface is more nuanced than Fe or Cu, where the host leverages its intricate means of manipulating Zn availability and distribution to limit the ability of the pathogen to access Zn while simultaneously ensuring sufficient Zn to support the immune response.

In addition to representing a distinct model system for studying nutritional immunity, we anticipated that the presence of functional Slc11a1 would affect the global immune response to infection, since Slc11a1 profoundly affects infection outcomes, namely resistance versus susceptibility. To critically compare infection-induced changes in the transcriptome, we identified an RNAseq study that was carried out in macrophages from C57BL/6 mice(47), with strong parallels to our conditions, and reanalyzed the data along with ours in an identical pipeline. We found that 4500 genes were differentially expressed in one study but not the other, with differences in expression at baseline as well as in response to infection. This is consistent with studies of mouse lines possessing one or two functional *slc11a1* alleles on a C57BL/6 background(30, 49). These mice experienced lower infection loads but still ultimately succumbed to *Salmonella* infection, implying differences between C57BL/6 mice and 129S6 mice that extend beyond Slc11a1. However, the magnitude of differential expression was unexpected, as it involved more than a third of all genes differentially expressed in our data. Annotation analysis of this differential gene list indicates that these cell types tend toward different fates in response to the stress of *Salmonella* infection, with 129S6 cells showing suppression of cell cycle genes but C57BL/6 cells showing upregulation of apoptotic genes.

In assessing the GSEA and GO term enrichment present in our data we find recurring themes of inflammation, hypoxia, apoptosis and the unfolded protein response (UPR), which are inextricably linked(36, 55). Both intracellular hypoxia and *Salmonella*-induced inflammation can result in apoptosis or the UPR, which is an attempt to regain cellular homeostasis under stress. When comparing live vs HK *Salmonella* conditions across time, we see enrichment of hypoxia-related genes along with enrichment of apoptosis 2 Hrs, which transitions to enrichment of UPR at 18 Hrs. This apparent transition in the cellular stress response over time demonstrates the ability of 129S6 macrophages to withstand *Salmonella* infection. In contrast, infection of C57BL/6 macrophages showed marked enrichment of apoptosis genes at 18 Hrs. This finding is supported by multiple previous studies which found that *Salmonella* induces apoptosis in Slc11a1 nonfunctional macrophages(39, 56).

Macrophages experience a range of phenotypes, from inflammatory and cytotoxic (termed M1 polarization) to anti-inflammatory and wound healing (M2 polarization), depending on their milieu. These states are fluid, differing in gene expression and metabolism, and are heterogeneous across *in vivo* cell populations(36). The bactericidal mechanisms available to M1 macrophages enable them to kill internalized pathogens efficiently. The lack of these mechanisms in M2-like macrophages make them a more permissive niche for facultative intracellular bacteria like *Salmonella*. Although a number of previous studies have tried to identify individual genes that distinguish M1 and M2, for example: *arg1* vs *nos2*(57) or *egr2* vs *CD38*(58), the increasing availability of whole transcriptome sequencing makes it possible to define a more comprehensive genetic signature of M1 and M2. Single cell RNAseq by Saliba et al(31) enabled identification of gene clusters related to M1 and M2 and how these gene clusters correlated with infection (naïve macrophages, bystanders, macrophages with growing bacteria and macrophages with non-growing bacteria). Dual RNAseq of both the pathogen and host by Stapels et al(47) further enables correlation of the M1 and M2 gene clusters within the host with expression of *Salmonella* virulence genes. Indeed, this analysis revealed that in C57BL/6 macrophages, the M2 polarization correlates with increased SPI2 expression, suggesting that *Salmonella* activates virulence mechanisms in this niche.

Examination of M1 and M2 gene clusters further reinforces that 129S6 macrophages represent a distinct intracellular niche with respect to immune response to infection. As shown in Fig 2, polarization genes follow a temporal pattern of expression, with the majority of M1 related genes strongly upregulated 2 Hrs post infection but sharply downregulated by 18 Hrs. On the other hand we see slower time-dependent activation of M2 related genes. This observation is supported at 2 Hrs by enrichment in glycolysis, the metabolic state of inflammatory or hypoxic macrophages, and at 18 Hrs by enrichment in oxidative phosphorylation and fatty acid metabolism, which are the metabolic markers of M2 macrophages(36, 55). Importantly, at each time point we see little difference in M1/M2 associated genes between live and HK *Salmonella* conditions, especially when normalized to C57BL/6 data **(Supplementary Figure S7)**. This is in contrast to findings in Slc11a1 non-functional macrophages which indicate that dividing *Salmonella* may actively induce M2 polarization to enhance bacterial survival, and that macrophages exposed to but not infected by *Salmonella* maintain an M1 phenotype late in infection(31, 47). While our work is at a population level and therefore contains macrophage and *Salmonella* heterogeneity, exposure to HK Salmonella does not lead to retention of an M1 phenotype at 18 Hrs, nor does it seem that 129S6 cells are induced to M2 by live *Salmonella*.

Our study recapitulates the ability of 129S6 macrophages to fight infection at all time points, which intriguingly occurs regardless of the expression profile of M1 or M2 genes. At 2 Hrs, 70% of initially infected macrophages had cleared the infection, which correlated with the transcriptional induction of M1 polarization. However, despite the shift toward an M2-like transcriptional profile at 18 Hrs, the rate of bacterial clearance increased slightly over time to 74%. Concurrently the proportion of infected macrophages showing evidence of bacterial replication also increased over time, from 8% at 10 and 18 Hrs to 24% by 24 Hrs. This high clearance rate and relatively low replication rate resulted in only 3% of initially infected macrophages harboring dividing bacteria at 24 Hrs **(Supplementary Dataset S5)**. This is in contrast to studies of Nramp1^+/+^ 129X1 mice(59) and Slc11a1 nonfunctional macrophages which report that replicating *Salmonella* are preferentially found in M2 cells(31,35,47). It is also notable that 129S6 macrophages surviving to 24 Hrs while harboring dividing *Salmonella* are not allowing high division rates, as indicated by only modest decreases in the inducible fluorescent signal (Fig 9), whereas Slc11a1 nonfunctional macrophages experience high levels of *Salmonella* division at 24 Hrs(35, 46). It is important to note that Zn was a key factor in both clearance and bacterial replication processes, as reduced zinc availability correlated with attenuation of both.

This study reveals that 129S6 macrophages differs significantly from other model systems of *Salmonella* infection which possess a nonfunctional Slc11a1. First, 129S6 macrophages show a robust remodeling of metal homeostasis and expression of metal-dependent enzymes, that indicate the nutritional immunity response within the host differs in the presence of functional Slc11a1. Second, remodeling of Zn regulatory proteins leads to an increase in cytosolic Zn and a likely restriction of Zn for the pathogen. This multi-pronged approach ensures sufficient cytosolic Zn to fight infection and promote bacterial clearance, while still limiting Zn availability to fight against bacterial replication. Third, transcription of M2-related genes doesn’t correlate exclusively with live *Salmonella* exposure or with a lack of bactericidal capacity, suggesting that in this model system the M2 polarization does not promote bacterial survival. Rather, activation of M1-related genes early and M2-related genes at later time points correlates with the switch from expression of apoptosis genes to UPR genes and suggests a choreographed response to fight infection without killing the host cell. Together, these imply that 129S6 macrophages do not conform well to the M1/M2 expression dichotomy derived largely from Nramp1 nonfunctional macrophages and mice. Instead, they appear to present a mixed phenotype that effectively uses zinc to quell *Salmonella* infection.

## Materials and Methods

### Ethics statement

This study was approved by the Institutional Animal Care and Use Committee at University of Colorado Boulder. The animal subjects plan protocol number is 2547.

### Monocyte extraction from bone marrow

Animal work followed protocol 2547, approved by University of Colorado IACUC. 8-12 week-old 129S6 female mice (Taconic Laboratories) were euthanized by CO_2_ inhalation according to IACUC guidelines, followed by cervical dislocation. Femur, tibia and humerus bones were extracted, then scraped and flushed with ice cold phosphate buffered saline (PBS). Approximately 1 mL of PBS was used per bone. PBS with marrow cells was passed through a 70 micron nylon mesh Falcon^TM^ Cell Strainer (Corning^TM^ 352350), then overlaid onto an equal volume of Histopaque-1083 (Sigma-Aldrich) in centrifuge tubes. Tubes were centrifuged at 500 x G for 30 min and allowed to slow with no brake. Monocytes appear as a fuzzy layer at the fluid interface. These were removed to clean tubes and washed twice with 14 mL PBS. After the final wash monocytes were resuspended in a small amount of macrophage growth media and counted with a hemocytometer. Growth media consisted of DMEM (Sigma-Aldrich) supplemented with 20 % fetal bovine serum (FBS), 2 mM L-glutamine, 1 mM sodium pyruvate, penicillin/streptomycin (50 IU/mL penicillin and 50 μg/mL streptomycin) and 10 pg/μL recombinant murine macrophage colony stimulating factor (PeproTech, Inc.).

### Primary cell culture and infection

Monocytes were plated into 12-well plates at a density of 200,000 cells per well. 2 mL macrophage growth media was added to each well and plates were incubated at 37 °C and 5% CO_2_ for 6 days. Media was refreshed 3 days after plating. Wild type *Salmonella* Typhimurium SL1344 was cultured to stationary phase in LB with [antibiotics] at 37 °C with aeration. Bacteria were washed with PBS and opsonized for 30 minutes at room temperature in a 1:1 solution of mouse serum (Sigma) and cell culture media (Gibco). Bacteria were then pelleted at 13,000 x G for 1 minute and resuspended in PBS. Half of the bacteria was heat killed at 60° C for 3 minutes, then placed on ice. One well of macrophages was scraped and counted, and bacteria were diluted in macrophage media without antibiotics to an MOI of 30. Plated macrophages were washed 3 times with PBS, then treated with antibiotic free macrophage media containing live, heat killed, or no bacteria. 30 minutes later bacterial media was removed, cells were washed 3x with PBS, and macrophage media containing 100 g/mL gentamicin was added. 90 minutes later media was again removed. RNEasy lysis buffer was added directly to wells designated as 2 Hr samples. Macrophage media with 10 μg/mL gentamicin was added to wells designated as 18 Hr samples. Lysed samples were collected and frozen at −20 °C. 18 Hrs post *Salmonella* exposure media was removed from remaining wells, lysis buffer was added, and cell lysates were frozen at −20 °C. To examine both early and late immune responses, macrophages were lysed at 2 or 18 hours post bacterial exposure. Control samples were lysed at the two-hour timepoint.

### RNA extraction and sequencing

RNA extraction was done on all samples at once, after one freeze-thaw cycle. An RNEasy kit (Qiagen) was used, and DNAse I treatment was done on the column. RNA integrity was checked via Tape Station, and library prep was done with an Illumina TruSeq LT kit which included polyA selection. Paired end 75 base sequencing was done on a NextSeq 2.1.0 Illumina machine. Tape station run, library prep and sequencing were performed by the JSCBB sequencing core.

### Data analysis pipeline

R 3.3.0 on the JSCBB computing core was used for analysis of raw data. Data quality was assessed with fastqc (0.11.2) and the first 10 bases were trimmed with Trimmomatic (0.36). Reads were mapped with TopHat(2.0.6)/Samtools (0.1.18)/bowtie2 (2.0.2) using --b2-very-sensitive, fr-firststrand settings. Read counting was completed with EdgeR Subread featureCounts (1.6.0) using the mm10 gtf file from UCSC (July 2015 version). MetaFeature mapping was paired end, with both ends mapped, and no MultiOverlap allowed. R 4.0.2 was used for all remaining analysis. A principal component analysis (PCA) of global RNA expression was performed using EdgeR. Analysis of differential expression was done with DESeq2 (1.18.1) but the nature of this dataset violates the DESeq2 assumption that most genes do not change expression between conditions. Therefore, all genes with differential expression padj < 0.05 in binary comparisons were compiled and excluded from the final dds normalization using the controlGenes parameter. A log ratio test on the correctly normalized DESeq dataset matrix (dds) with the following function: DESeq(dds, test=“LRT”, reduced = ∼ 1), and resulted in a list of globally differentially expressed genes. This dataset (p < 0.01) was further filtered with two Transcripts Per Million (TPM) requirements. Inclusion in the final analysis indicates that a gene has an overall mean TPM > 5, and at least one condition with a mean TPM > 10. The 7766 genes meeting these criteria were fed into DEGReport R package(1.24.1) for rlog transformation, which produced normalized counts, and expression clustering, which produced Z-scores. Expression clustering was done with the following function: degPatterns(cluster_rlog, metadata = metadata, time = “Treatment”).Heatmap gene dendrograms were constructed using hierarchical clustering package hclust and ggdendro (0.1.22), and this gene order was used to construct global and cluster heatmaps with ggplot (Tidyverse R package 1.3.0). The median value for each plot was set as white. DAVID annotation enrichment categories were used to assess the makeup of DEGReport clusters. This analysis included functional categories, Gene Ontology, Biocarta and Kegg pathways, and protein domains.

In brief, DESeq2 binary comparisons between conditions were fed into GSEA(60) to validate GO term enrichment and assess transcription targets. More specifically, adjusted p-value ranked lists were created from DESeq2 differential expression data sets comparing 2A vs. 2HK or 18A vs. 18HK. For each gene in a given data set, a rank was determined from the padj value (false discovery rate) and the direction of fold change value (up or down regulation). Rank = −log(padj) * sign(fold change). All genes with a valid padj value were included. GSEA was used to compare these ranked lists to validated gene sets. These were: Canonical Pathways, Hallmark gene sets, GO_Biological Processes, GO_Molecular Functions, and Transcription Factor Targets. Because GSEA’s default is human gene sets, the mouse genome was collapsed and remapped using the Mouse Gene Symbol Remapping Human Orthologs MsigDB.v7.2.chip. Gene set permutations were used in this analysis, which is a less stringent assessment of significance, and produces more false positive results. Therefore, a cutoff of q < 0.05 was applied to gene set results.

### Comparison of current study with Staples et al, Science, 2018

FASTQ files (GEO accession GSE104785) were downloaded and run through the above pipeline with the following alterations: Trimmomatic was used to remove Illumina adaptors, and alterations were made to mapping and counting based on the unstranded nature of the data. This differs from the pipeline of Staples et al. in that their data was mapped to the *Salmonella* genome prior to the mouse mm10 genome. Additionally, their read counts were performed with HTSeq (version 0.6.1) instead of FeatureCounts. To normalize our data with this previously published dataset, DESeq2 was run on the combined data with the experimental design ∼Treatment, and normalized counts were calculated. PCA and dendrogram of the Staples dataset were created and compared with their published heatmap to generally validate our sequencing results (**Supplementary Figure S8**).

To assess differences in metal homeostasis genes as comprehensively as possible, mouse specific lists of genes involved in the transport and homeostasis of metal ions were compiled from the Uniprot database. Separate lists were compiled for Zn, Fe, and Cu, and each list was compared against the normalized counts of the aggregated sequencing studies. Only genes expressed above background were used for further analysis.

### Measurement of cytosolic Zn and Imaging analysis

For quantification of cytosolic labile Zn in primary macrophages, calibration experiments were performed at the indicated times post infection on a Nikon Ti-E widefield fluorescence microscope equipped with Nikon elements software, Ti-E perfect focus system, an iXon3 EMCCD camera (Andor), mercury arc lamp, and YFP FRET (434/16 excitation, 458 dichroic, 535/20 emission), CFP (434/16 excitation, 458 dichroic, 470/24 emission), and YFP (495/10 excitation, 515 dichroic, 535/20 emission) filter sets. External excitation and emission filter wheels were controlled by a Lambda 10-3 filter changer (Sutter Instruments), while dichroic mirrors were placed on cubes in the dichroic turret. Channel EM gain and exposure settings for both YFP FRET and CFP were set to be the same. Images were collected using a 60X oil objective (NA 1.40), 200 ms exposure time, EM gain 1 MHz at 16-bit readout mode with an EM gain multiplier of 200, and a neutral density filter with 25% light transmission. Sensor expression level was controlled by selecting cells with YFP intensities between 4,000-15,000 fluorescence units under these conditions. 8 fields of view were collected using multipoint acquisition mode and the Perfect Focus System (PFS) set to ‘ON’ in between points. Cells were maintained at 37°C and 5% CO2 in a LiveCell^TM^ environment chamber (Pathology Devices) during the experiments. Images were collected every minute during R_Resting_ and R_min_ calibration acquisition phases and every 30 seconds during the R_max_ calibration phase.

Fresh calibration solutions were prepared the day of the experiment and include a 2X solution of R_min_ buffer (50µM TPA in PO_4_^3-^-free HHBSS) for minimum FRET ratio collection and a 2X solution of R_max_ buffer (0.001% saponin + 0.75 μM pyrithione + 23.8nM buffered Zn in PO_4_^3-^-, Ca^2+^-, Mg^2+^-free HHBSS) for collecting the maximum FRET ratio. The resting FRET ratio of the sensor was collected for 10 mins prior to calibration to ensure a stable signal. After 10 min of imaging 50 µM TPA was added to the dish to collect the minimum FRET ratio of the sensor. Once a stable signal had been achieved cells were then washed with phosphate, calcium, and magnesium-free HEPES-buffered HBSS, pH = 7.4 to remove the chelate and then treated with pyrithione and Zn with 0.001% (w/v) saponin.

All imaging data were analyzed in MATLAB (Mathworks). Images were background corrected by subtracting a local background intensity from each pixel grouped in a certain region of the image. Regions of interest were generated by using a segmentation algorithm that segments the image based on the fluorescence intensity of the FRET ratio channel to obtain single cell traces. FRET ratios for each cell trace were calculated by dividing the background-corrected YFP FRET intensity by the background-corrected CFP intensity. The resting, minimum, and maximum FRET ratios for each cell were used to calculate the fractional saturation, the dynamic range, and the reported [Zn] from the sensor in each cell. Resting [Zn] is calculated by[*Zn*^2+^] = 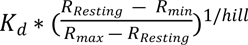 where the K_d_ = 230 pM and hill = 0.53 for NES-ZapCV2. The dynamic range of the sensor can be impacted by both over expression and under expression of the sensor which has a negative impact on the fidelity of the apparent [Zn]. For this reason, cells were excluded from the analysis if they fell outside of an acceptable dynamic range, 1.6 – 2.3 for the NES-ZapCV2 sensor.

### Flow cytometry study of infection outcome with varied zinc availability

Four different media conditions were used for flow cytometry. Standard macrophage growth media (DMEM + 20% FBS) was found to contain 25 μM Zn by ICP-MS and was used as the baseline. The other three medias used chelex-100-treated FBS. Briefly, FBS was treated with Chelex 100 resin (Sigma-Aldrich, St. Louis, MO, USA) for 5 hours, stirring at 4 °C, followed by sterile filtration using a 0.22 µm PES membrane filter. This chelex-treated FBS was used to prepare the baseline Chelex media, which contained DMEM + 20% FBS + L-glutamine (2 mM) + sodium pyruvate (1 mM). Zn replete media was made by adding 30 µM ZnCl_2_ (Sigma-Aldrich, St. Louis, MO, USA) to Chelex media. Media with essentially no Zn was made by adding 3 µM 2-{[Bis(2-pyridinylmethyl)amino]ethylamino}benzenesulfonic (ZX1, an extracellular Zn chelator(61)) (Strem Chemicals, Inc.) to Chelex media. ICP-MS measurements were carried out as described previously(62) to quantify the metal content of the media. While Chelex treatment depleted Zn, Ca, and Fe, the Zn replete, Chelexed media, and Zn deficient media are identical except for the amount of zinc present in the media (34 μM, 4 μM, no Zn, respectively).

For analysis of intracellular bacterial replication, monocytes from three mice were grown and differentiated to macrophages in 6 well plates. Six days later, *Salmonella* expressing pDiGc and induced with arabinose were grown to stationary phase with aeration. Macrophages were infected with *Salmonella* at a multiplicity of infection of 30, and infection proceeded for 45 min. At this point *Salmonella* media was removed, cells were washed 3x in PBS, and initial inoculum cells were collected by scraping with a nylon cell lifter (Corning™ C3008) and homogenized by pipetting gently with a P1000. Homogenized cells were fixed in a gentle fixative for preserving fluorescent protein fluorescence (1 % PFA and 1 % sucrose) for 15 min and then washed and resuspended in PBS and chilled at 4 °C. Media in all other samples was changed to experimental conditions which included 10 μg/mL gentamicin. Three wells were treated with each media, for each time point. At 2, 10, 18 and 24 Hrs post infection, cells were rinsed, lifted, fixed and chilled as above. Samples were analyzed on a BD FACSCelesta™ (BD Bioscences) collecting forward scatter area and width, side scatter area and width, 488 nm excitation 530/30 nm emission, and 561 nm excitation and 585/15 nm emission. Data were analyzed using FlowJo 10.5.3 software (FlowJo LLC).

The cell gating hierarchy was set as Single cells > GFP positive cells > Cells containing replicated bacteria. 25-30,000 single cells were collected per sample. Single cells were determined first by forward scatter area versus side scatter area then by side scatter width versus side scatter area. Non-fluorescent uninfected cells were used to set the gate for GFP positive cells. A ratio of the 488 nm channel to the 561 nm channel was taken by dividing 488 nm intensity by 561 nm intensity. Samples collected at 2 Hrs post infection were used as the “initial inoculum” to determine the fluorescence intensities for cells infected with bacteria that have not undergone replication. Cells containing replicated bacteria were gated as having a 488 nm:561 nm ratio above the initial inoculum.

## Acknowledgements

We would like to thank Dr. Lynn Sanford for technical assistance developing code for analyzing RNAseq data. We acknowledge the BioFrontiers Institute Advanced Light Microscopy Core, where spinning disc confocal microscopy was performed on a Nikon Ti-E microscope supported by the BioFrontiers Institute and the Howard Hughes Medical Institute. We would like to acknowledge the University of Colorado BioFrontiers Institute Next-Gen Sequencing Core Facility, which performed the Illumina sequencing and library construction, the BioFrontiers high-performance computing resources (NIH 1S10OD012300) supported by BioFrontiers’ IT, and the University of Colorado Biochemistry Cell Culture Core Facility for providing resources and support in culturing.

## Data availability

All raw next-generation sequencing data files and processed data files used to draw conclusions are available at the Gene Expression Omnibus, data series GSE166642.

